# The Developing Human Connectome Project: a Minimal Processing Pipeline for Neonatal Cortical Surface Reconstruction

**DOI:** 10.1101/125526

**Authors:** Antonios Makropoulos, Emma C. Robinson, Andreas Schuh, Robert Wright, Sean Fitzgibbon, Jelena Bozek, Serena J. Counsell, Johannes Steinweg, Katy Vecchiato, Jonathan Passerat-Palmbach, Gregor Lenz, Filippo Mortari, Tencho Tenev, Eugene P. Duff, Matteo Bastiani, Lucilio Cordero-Grande, Emer Hughes, Nora Tusor, Jacques-Donald Tournier, Jana Hutter, Anthony N. Price, Rui Pedro A. G. Teixeira, Maria Murgasova, Suresh Victor, Christopher Kelly, Mary A. Rutherford, Stephen M. Smith, A. David Edwards, Joseph V. Hajnal, Mark Jenkinson, Daniel Rueckert

## Abstract

The Developing Human Connectome Project (dHCP) seeks to create the first 4-dimensional connectome of early life. Understanding this connectome in detail may provide insights into normal as well as abnormal patterns of brain development. Following established best practices adopted by the WU-MINN Human Connectome Project (HCP), and pioneered by FreeSurfer, the project utilises cortical surface-based processing pipelines. In this paper, we propose a fully automated processing pipeline for the structural Magnetic Resonance Imaging (MRI) of the developing neonatal brain. This proposed pipeline consists of a refined framework for cortical and sub-cortical volume segmentation, cortical surface extraction, and cortical surface inflation, which has been specifically designed to address considerable differences between adult and neonatal brains, as imaged using MRI. Using the proposed pipeline our results demonstrate that images collected from 465 subjects ranging from 28 to 45 weeks post-menstrual age (PMA) can be processed fully automatically; generating cortical surface models that are topologically correct, and correspond well with manual evaluations of tissue boundaries in 85% of cases. Results improve on state-of-the-art neonatal tissue segmentation models and significant errors were found in only 2% of cases, where these corresponded to subjects with high motion. Downstream, these surfaces will enhance comparisons of functional and diffusion MRI datasets, supporting the modelling of emerging patterns of brain connectivity.

## 1. Introduction

The period of rapid cortical expansion during fetal and early neonatal life is a crucial time over which the cortex transforms from a smooth sheet to a highly convoluted surface. During this time, the cellular foundations of our advanced cognitive abilities are mapped out, as connections start to form between distant regions (Ball et al., 2013b; Van Essen, 1997), myelinating, and later pruning, at different rates. Alongside the development of this neural infrastructure, functional brain activations start to be resolved (Doria et al., 2010), reflecting the development of cognition.

Much of what is currently known about the early human connectome has been learnt from models of preterm growth (Ball et al., 2013b; Counsell et al., 2013; Doria et al., 2010; Keunen et al., 2017). Whilst invaluable, it is known that early exposure to the extra-uterine environment has longterm implications (Ball et al., 2012, 2013a, 2017; Counsell et al., 2014; Hintz et al., 2015; Ullman et al., 2015). For this reason, the Developing Human Connectome Project (dHCP) seeks to image emerging brain connectivity for the first time in a large cohort of fetal and term-born neonates.

More broadly, the goals of the dHCP are to pioneer advances in structural, diffusion, and functional Magnetic Resonance Imaging (MRI), in order to collect high-quality imaging data for fetuses and (term/preterm-born) neonates, from both control and at-risk groups. Imaging sets will be supported by a database of clinical, behavioural, and genetic information, all made publicly available via an expandable and user-friendly informatics structure. The dataset will allow the community to explore the neurobiological mechanisms, and genetic and environmental influences, which underpin healthy cognitive development. Models of healthy development will provide a vital basis of comparison from which the effects of preterm birth, and neurological conditions such as cerebral palsy or autism, may become better understood.

The dHCP takes inspiration from the WU-MINN Human Connectome Project (HCP) (Van Essen et al., 2013). Now in its final stages, the HCP has pushed the boundaries of MRI based brain connectomics, collecting 1200 sets of healthy adult functional and structural connectomes, at high spatial and temporal resolution. Data from this project has been used to generate refined maps of adult cortical organisation (Glasser and Van Essen, 2011), and improve understanding of how the functional connectome correlates with behaviour (Smith et al., 2015)

A core tenet, underlying the success of the HCP approach, has been the advocation of surface-based processing and analysis of brain MR images. This is grounded in the understanding that distances between functionally specialised areas on the convoluted cortical sheet are more neurobiologically meaningful, when represented on the 2D surface rather than in the 3D volume (Glasser et al., 2013). Surface-based processing therefore minimises the mixing of data from opposing sulcal banks or between tissue types. Further, surface-based registration approaches (Durrleman et al., 2009; Fischl et al., 1999c; Lombaert et al., 2013; Robinson et al., 2014; Wright et al., 2015; Yeo et al., 2010) improve the alignment of cortical folds and areal features.

Unfortunately, modelling cortical connectome structure in neonates and fetuses is particularly challenging. MRI, especially functional and diffusion protocols, is highly sensitive to head motion during scanning. This is a particular issue for the dHCP, where the goal is to image un-sedated neonatal, and free-moving fetal subjects. Therefore, correcting for this has required the development of advanced scanning protocols and motion correction schemes (Cordero-Grande et al., 2017; Hughes et al., 2016; Kuklisova-Murgasova et al., 2012).

Outside of the challenges facing acquisition and reconstruction, the properties of neonatal and fetal MRI differ significantly from that of adult data. Specifically, baby and adult brains differ vastly in terms of size, with the fetal and neonatal brain covering a volume in the range of 100-600 millilitres in contrast to an average adult brain volume of more than 1 litre (Allen et al., 2002; Orasanu et al., 2014; Makropoulos et al., 2016). Furthermore, the perinatal brain develops rapidly, which results in vast changes in scale and appearance of the brain scanned at different weeks. This, together with the fact that scanning times must be limited for the well-being and comfort of mother and baby, means that spatial and temporal resolution of the resulting images are reduced relative to adults. Furthermore, immature myelination of the white matter in neonatal and fetal brains results in inversion of MRI contrast when compared to adult brain scans (Prastawa et al., 2005). This requires image processing to be performed on T2-weighted rather than T1-weighted structural MRI.

Combined, these vast differences in image properties considerably limit the translation of conventional adult methods for image processing to fetal and neonatal cohorts. In particular, the popular FreeSurfer framework (Fischl, 2012), utilised within the HCP pipelines (Glasser et al., 2013), fails on neonatal data as it relies solely on fitting surfaces to intensity-based tissue segmentation masks (Dale et al., 1999). These pipelines have been optimised to work with adult MRI data, and are not compatible with neonatal image intensity distributions, which are significantly different and vary drastically within different weeks of development. For this reason, simple adoption of the adult HCP processing pipelines has not been possible.

Instead, this paper presents a refined surface extraction and inflation pipeline, that will accompany the first data and software release of the dHCP. This proposed framework builds upon a legacy of advances in neonatal image processing. This includes the development of specialised tools for tissue segmentation that address the difficulties in resolving tissue boundaries blurred through the presence of low resolution and partial volume. A variety of techniques have been proposed for tissue segmentation of the neonatal brain in recent years: unsupervised techniques (Gui et al., 2012), atlas fusion techniques (Weisenfeld and Warfield, 2009; Gousias et al., 2013; Kim et al., 2016), parametric techniques (Prastawa et al., 2005; Song et al., 2007; Xue et al., 2007; Shi et al., 2010; Cardoso et al., 2013; Makropoulos et al., 2012; Wang et al., 2012; Wu and Avants, 2012; Beare et al., 2016; Liu et al., 2016), classification techniques (Anbeek et al., 2008; Srhoj-Egekher et al., 2012; Chiţa et al., 2013; Wang et al., 2015; Sanroma et al., 2016; Moeskops et al., 2016) and deformable models (Wang et al., 2011; Dai et al., 2013; Wang et al., 2013, 2014). A review of neonatal segmentation methods can be found in Devi et al. (2015); Makropoulos et al. (2017). The majority of these techniques have been applied to images with a lower resolution than those acquired within the dHCP, and typically to images of preterm-born subjects.

Once segmentations are extracted, surface mesh modelling approaches are, to an extent, agnostic of the origin of the data; allowing, in principle, the application of a wide variety of cortical mesh modelling approaches to neonatal data, including those provided within the FreeSurfer (Dale et al., 1999; Fischl, 2012), BrainSuite (Shattuck and Leahy, 2002), BrainVISA (Riviere et al., 2009), and CIVET (MacDonald et al., 2000; Kim et al., 2005, 2016) packages. In general, these methods fit surfaces to boundaries of tissue segmentation masks, which, allowing for some need for topological correction, relies on the accuracy of the segmentation. In neonatal imaging data, the use of T2 images however leads to segmentation errors not seen in adult data. This is the misclassification of CSF as white matter, caused by the fact that CSF and white matter appear bright in neonatal T2 images, whereas in adult T1 data white matter is bright and CSF is dark. If not fully accounted for during segmentation, these errors will be propagated through to surface reconstruction (Xue et al., 2007).

In what follows, we present a summary of our proposed pipeline. This brings together existing tools for neonatal segmentation (refined to minimise the propagation of misclassification errors through to surface extraction) with new tools for cortical extraction that combine information from segmentation masks and T2-weighted image intensities, in order to compensate for the effects of partial volume and improve correspondence with true tissue boundaries. Re-implementations of existing tools for surface inflation and projection to the sphere are provided to minimise software overhead for users.

The processing steps are as follows: 1) acquisition and reconstruction of T1 and T2 images (Cordero-Grande et al., 2017; Hughes et al., 2016; Kuklisova-Murgasova et al., 2012); 2) tissue segmentation and regional labelling (Makropoulos et al., 2012, 2014, 2016); 3) cortical white and pial surface extraction (Schuh et al., 2017); 4) inflation and projection to a sphere (for use with spherical alignment approaches) (Fischl et al., 1999a; Elad et al., 2005); and 5) definition of cortical feature descriptors, including descriptors of cortical geometry and myelination (Glasser et al., 2013). Manual quality control is performed by two independent expert raters to assess the quality of the acquired images, segmentations, and reconstructed cortical surfaces. Assessment of these data sets shows, that with very few exceptions (3%) the protocol is able to extract cortical surfaces which fit closely the expectations for observed anatomy.

## 2. Project Overview

The goal of the dHCP is to create a dynamic map of human brain connectivity from 20 to 45 weeks post-conceptional age (PMA) from healthy, term-born neonates, infants born prematurely (prior to 36 weeks PMA), and fetuses. The infants are being scanned using optimised protocols for structural (T1 and T2-weighted) images, resting state functional MRI (fMRI), and multi-shell High Angular Resolution Diffusion Imaging (HARDI). Imaging data will be combined with genetic, cognitive, and environmental information in order to aid understanding of the developing human brain, and to give crucial insight into brain vulnerability and disease development. Novel image analysis and modelling tools will be developed and integrated into HCP inspired surface-processing pipelines in order to extract structural and functional connectivity maps. All data and supporting software will be made publicly available within an expandable, future-proof informatics structure, which will provide the research community with a user-friendly environment for hypothesis-based studies, and allow continual ongoing addition of new data.

## 3. The Neonatal Structural Pipeline

The first stage of the project has been to optimise acquisition protocols and collect data for the neonatal cohort (Cordero-Grande et al., 2017; Hughes et al., 2016; Kuklisova-Murgasova et al., 2012). The methods in this paper therefore reflect neonatal structural processing protocols, and are designed to accompany the first data release of neonatal subjects.

The workflow of the neonatal processing pipeline is summarised in Fig. 1. The motion-corrected, reconstructed T2-weighted image is first bias-corrected and brain-extracted. Following this, the brain image is segmented into different tissue types (CSF: cerebrospinal fluid, WM: white matter, cGM: cortical grey matter, and GM: subcortical grey matter) using the Draw-EM algorithm (based on Makropoulos et al. (2014) section 3.2) (B). Next, white matter masks are split along the mid-line between the hemispheres and filled to represent binary masks for each hemisphere, and topologically correct white matter surfaces are fit first to the grey-white tissue boundary and then the grey-white interface of the MR intensities (section 3.3) (C). Pial surfaces are generated by expanding each white matter mesh towards the grey-CSF interface (D), and midthickness surfaces are generated halfway between the pial and the white, by taking the Euclidean mean of corresponding inner- and outer-cortical vertex coordinates (E). The cortical thickness is estimated based on the Euclidean distance between the white and pial surface. In a separate process, inflated surfaces are generated through expansion-based smoothing of the white surface (F). This acts as initialisation to a Multi-Dimensional Scaling (MDS, section 3.4) scheme that projects all points onto a sphere, whilst preserving relative distances between neighbouring points. Finally, surface geometry and myelo-structure are summarised through a series of surface feature maps. These include: G) maps of mean surface curvature, estimated from the white surface; H) maps of mean convexity/concavity (sulcal depth); estimated during inflation; and I) maps of cortical myelination; estimated from the ratio of T1- and T2-weighted intensities projected onto the surface. Prior to myelin estimation, the motion-corrected reconstructed T1 image is rigidly registered to the T2 image.

**Figure 1:**
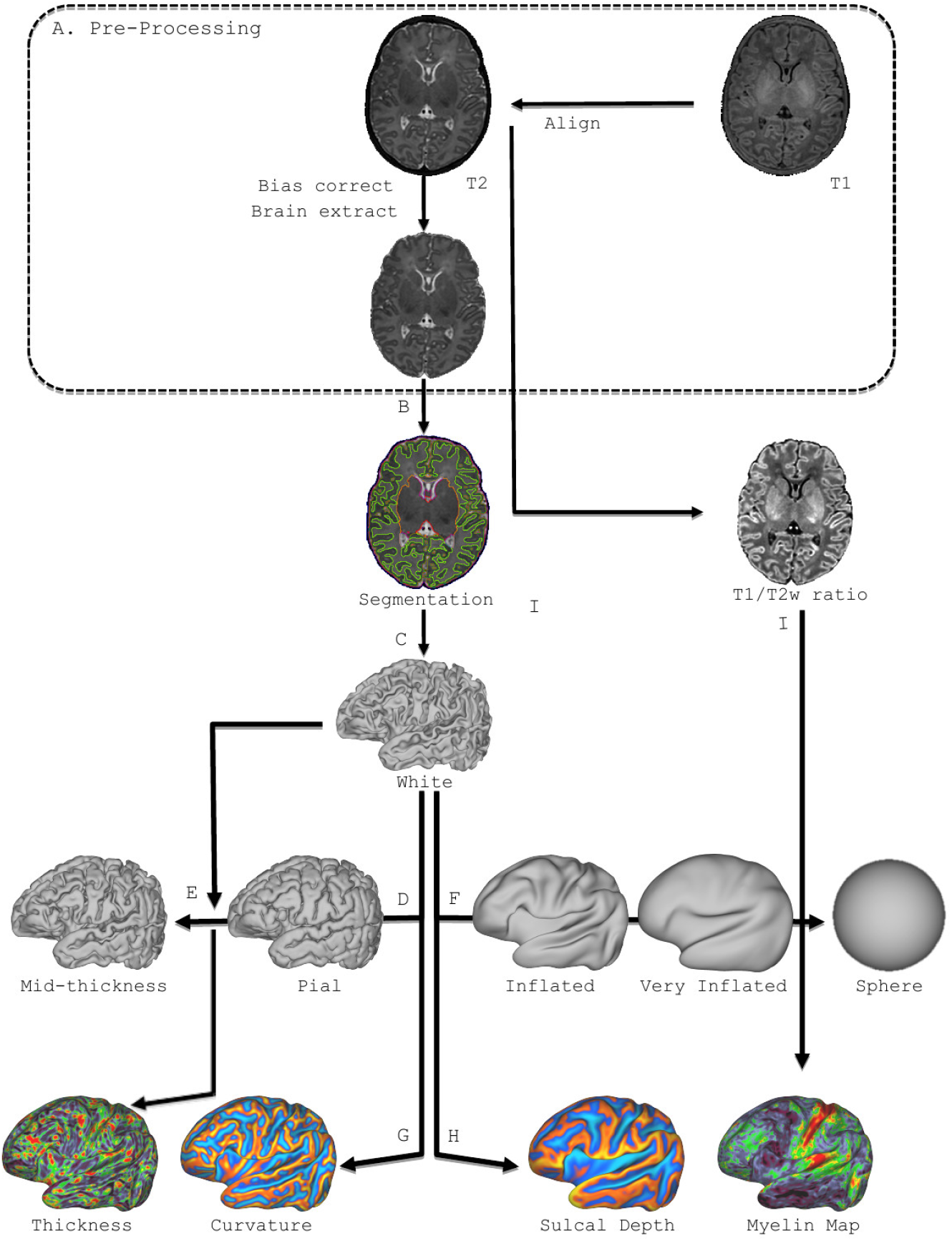
Structural pipeline steps: A) Pre-processing; B) Tissue segmentation (shown as contour map); C) White-matter mesh extraction; D) Expansion of white surface to fit the pial surface; E) Fitting of midthickness surface midway between white and pial and estimation of thickness; F) Inflation of white surface to fit the inflated surface and projection onto a sphere; G) Estimation of surface curvature from the white surface; H) Estimation of sulcal depth maps (mean convexity/concavity); I) Estimation of T1/T2 myelin maps;

Following adult frameworks, proposed for FreeSurfer (Fischl et al., 1999b), and refined by the Human Connectome Project (Glasser et al., 2013), all computed surfaces of each subject (white, pial, midthickness, inflated, spherical) have one-to-one point correspondence between vertices. This ensures that for each individual brain the same vertex index represents the same relative point on the anatomy for all these surfaces. In other words, two corresponding points on the white and pial surface would approximate the endpoints of where a column of neurons through the cortex would travel. This simplifies downstream processing, and facilitates straightforward visualisation of features across multiple surface views.

### 3.1. Acquisition

The data used in the paper were collected at St. Thomas Hospital, London, on a Philips 3T scanner using a 32 channel dedicated neonatal head coil (Hughes et al., 2016). To reduce the effects of motion, T2 images were obtained using a Turbo Spin Echo (TSE) sequence, acquired in two stacks of 2D slices (in sagittal and axial planes), using parameters: TR=12s, TE=156ms, SENSE factor 2.11 (axial) and 2.58 (sagittal) with overlapping slices (resolution (mm) 0.8 × 0.8 × 1.6). T1 images were acquired using an IR (Inversion Recovery) TSE sequence with the same resolution with TR=4.8s, TE=8.7ms, SENSE factor 2.26 (axial) and 2.66 (sagittal). Motion correction and super-resolution reconstruction techniques were employed combining Cordero-Grande et al. (2017); Kuklisova-Murgasova et al. (2012) resulting in isotropic volumes of resolution 0.5 × 0.5 × 0.5mm^3^. Subjects were not sedated, but imaged during natural sleep. All images were reviewed by an expert paediatric neuroradiologist and checked for possible abnormalities. In this paper we report results from 453 subjects that have been processed through the proposed pipeline. 38 subjects had repeat longitudinal scans resulting in a total 492 scans. T1 images were available for 411 out of the 492 scans. The distribution of ages at birth and ages at scan is shown in Fig. 2.

**Figure 2:**
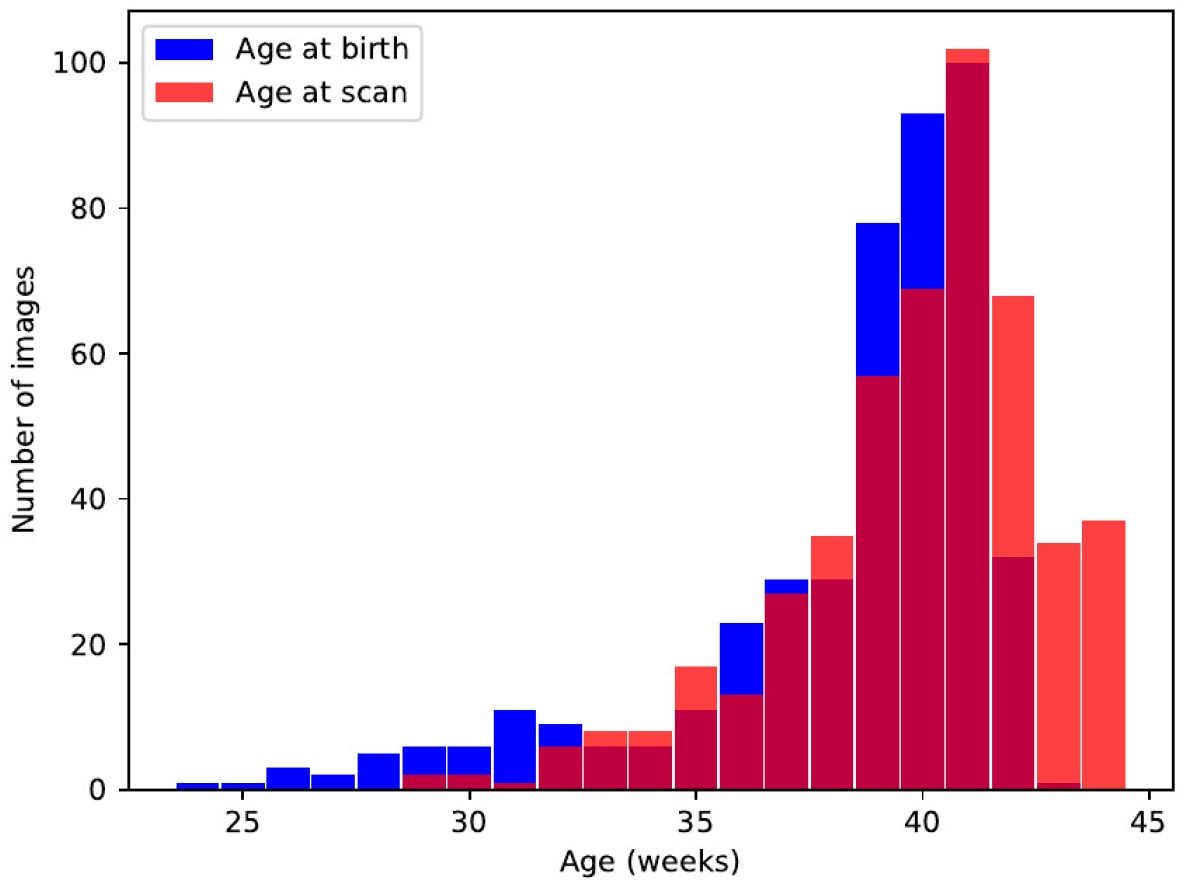
Distribution of age at birth and age at scan for the 453 processed subjects (492 scans).

### 3.2. Segmentation

#### 3.2.1. Atlases

The segmentation in the dHCP utilises the atlases manually annotated by Gousias et al. (2012) that divide the brain into 50 regions. Gousias et al. (2012) have manually labelled 20 subjects and provide T1 and T2 images accompanied by the label map for each subject^2^. 15 of the 20 subjects were born prematurely at a median age at birth of 29 weeks (range 26 – 35 weeks). These were scanned at term at a median age of 40 weeks (range 37 – 43 weeks), and had a median weight of 3.0 kg (range 2.0 – 4.0 kg) at the time of scan. The remaining 5 subjects were born at term at a median age of 41 weeks (range 39 – 45 weeks), and had a median weight of 4.0 kg (range 3.0 – 5.0 kg). We refer the reader to Gousias et al. (2012) for information on the protocol used for the labelling. In Makropoulos et al. (2014) we further subdivided the 50 regions to create a set of 87 regions as follows: the cortical regions, which contain both WM and cGM in the atlases, were split into their WM and cGM part; thalamus was split into the high intensity part and the low intensity part (mainly ventrolateral nuclei); CSF, background and unlabelled brain region (mainly internal capsule) were further added. Table 1 presents the 87 resulting regions.

**Table 1:** Tissue labels

| Tissues |  |
| --- | --- |
| 1 | Cerebrospinal fluid (CSF) |
| 2 | Cortical grey matter (cGM) |
| 3 | White matter (WM) |
| 4 | Background |
| 5 | Ventricle |
| 6 | Cerebellum |
| 7 | Deep Grey Matter (GM) |
| 8 | Brainstem |
| 9 | Hippocampus |

#### 3.2.2. Segmentation method

The T2 image data is segmented into 9 tissue classes (Table 1), using the Draw-EM ^3^ (Developing brain Region Annotation With Expectation-Maximization) algorithm. This is an open-source software for neonatal brain MRI segmentation based on the method proposed in Makropoulos et al. (2014).

**Table 2:**
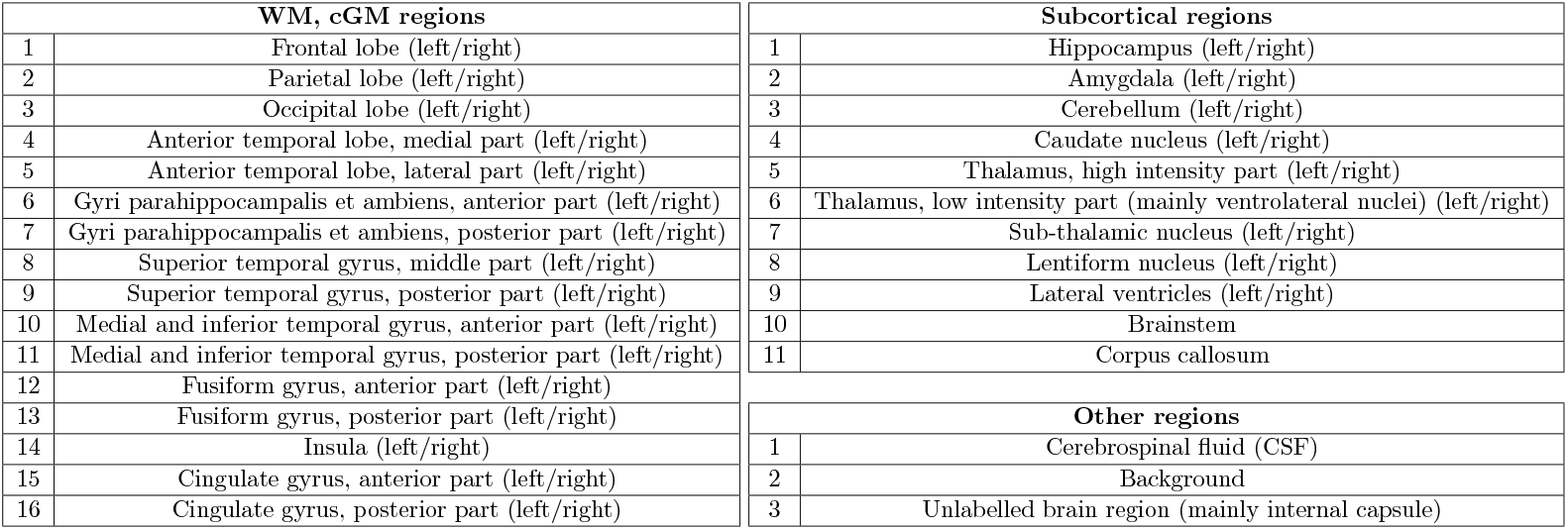
87 regional labels segmented with Draw-EM, based on the 20 atlases of (Gousias et al., 2012).

In summary, segmentation proceeds following brain-extraction, implemented using the Brain Extraction Tool (BET) (Smith, 2002), in order to remove non-brain tissue. BET has been configured to provide a liberal mask that always retains the brain tissue and CSF, and removes most of the skull (fractional intensity threshold=0.1). An accurate initial brain mask is not crucial for the Draw-EM segmentation, as it is based on an adaptive Expectation-Maximization algorithm. The brain mask is consequently refined from the Draw-EM tissue segmentation. Then images are corrected for intensity inhomogeneity with the N4 algorithm (Tustison et al., 2010).

Segmentation is performed with an Expectation-Maximization (EM) algorithm that includes a spatial prior term and an intensity model of the image, similar to the approach described in Van Leemput et al. (1999). The spatial prior probability of the different brain structures is defined based on multiple labelled atlases. These atlases are registered to the target image, and their labels are transformed and averaged according to the local similarity of each atlas (based on the local Mean Square Distance as proposed in Artaechevarria et al. (2009). The intensity model of the image is approximated with a Gaussian Mixture Model (GMM). Spatial dependencies between the brain structures are modelled with Markov Random Field (MRF) regularisation. The segmentation algorithm further includes CSF-WM partial volume correction to account for the similar intensity of CSF to the WM along the CSF-cGM boundary, and model averaging of EM and label fusion to limit the influence of intensity in the delineation of structures with very similar intensity profiles. The resulting segmentation contains 87 regional structures (see Table 1). These are merged to further produce the following files: tissue labels, left/right white surface masks and left/right pial surface masks.

The dHCP data differs significantly from previous neonatal cohorts collected by our group, in that the vast majority of the data in previous studies were acquired from prematurely-born neonates and typically at lower spatial resolutions. Thus Draw-EM includes several modifications with respect to Makropoulos et al. (2014) in order to maximise the benefits for downstream processing. These include: modelling of additional tissue classes to account for the presence of hyper/hypo intense pockets of white matter, which occur naturally in the maturing brain; together with improvements afforded by a multi-channel volumetric registration, which simultaneously registers subjects’ T2 images and their grey-matter tissue segmentation masks (obtained from a preliminary run of the segmentation algorithm - see below).

All major steps of the Draw-EM pipeline are summarised in Fig. 6. An example segmentation using Draw-EM is illustrated in Fig. 7. The importance of the modifications to the segmentation approach are demonstrated through Figures 3 to 5. Specifically, Fig. 3 demonstrates improvements to segmentation accuracy through modelling two additional tissue classes: 1) low and 2) high intensity WM. Here, the low-intensity WM class is included to account for myelinated WM that is often mislabelled for cGM in particular around the insula and the frontal lobe, and the high-intensity WM class is included to correct for WM hyper-intensities around the ventricular area that are often mislabelled as ventricles. Low-intensity and high-intensity WM classes are merged to WM after the segmentation has completed.

**Figure 3:**
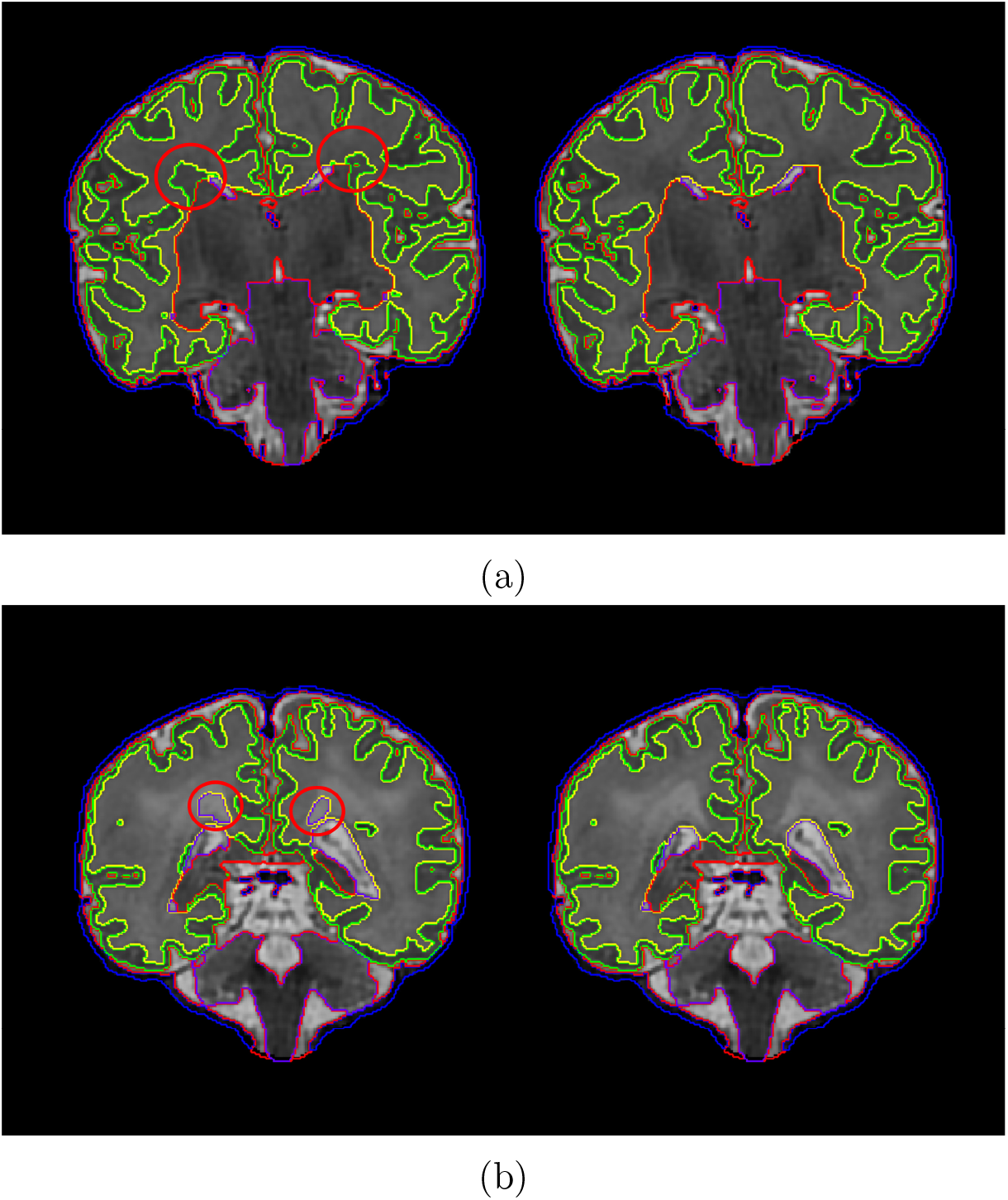
a) Segmentation of a subject T2 image without and with the inclusion of the low-intensity WM class (left and right image respectively). b) Segmentation of a subject T2 without and with the inclusion of the high-intensity WM class (left and right image respectively). Affected areas are noted with a red circle.

**Figure 4:**
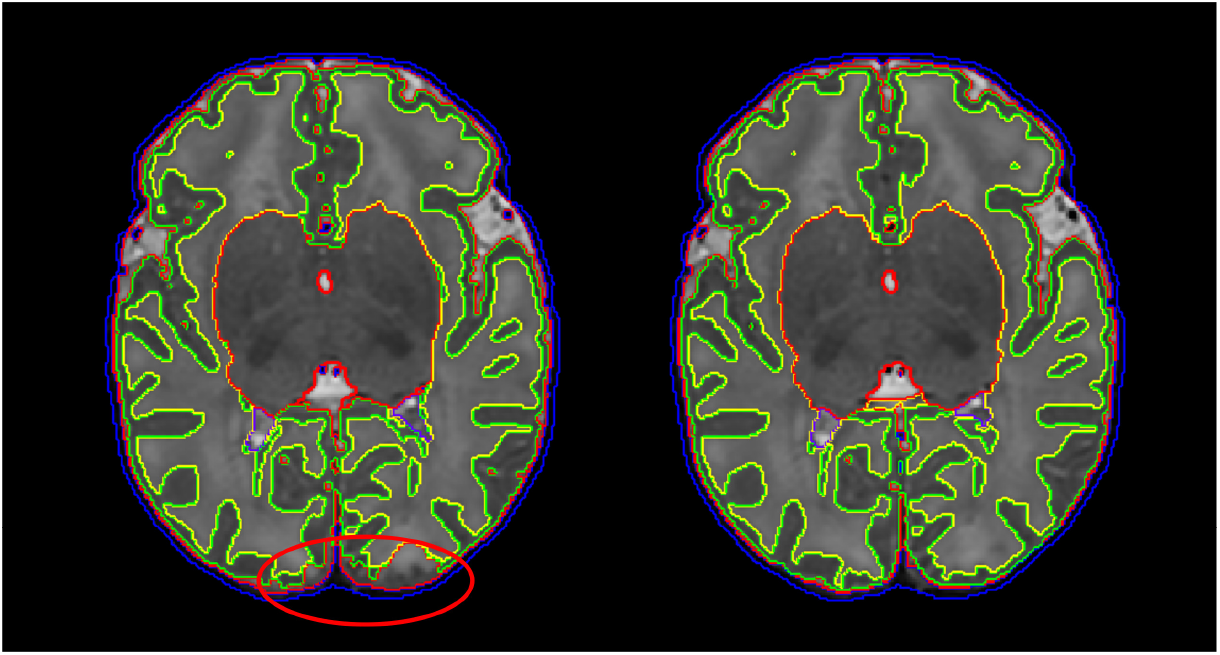
Segmentation of a subject T2 image based on the original preterm template Serag et al. (2012) and the trimmed template registration (right image). Affected areas are noted with a red circle.

**Figure 5:**
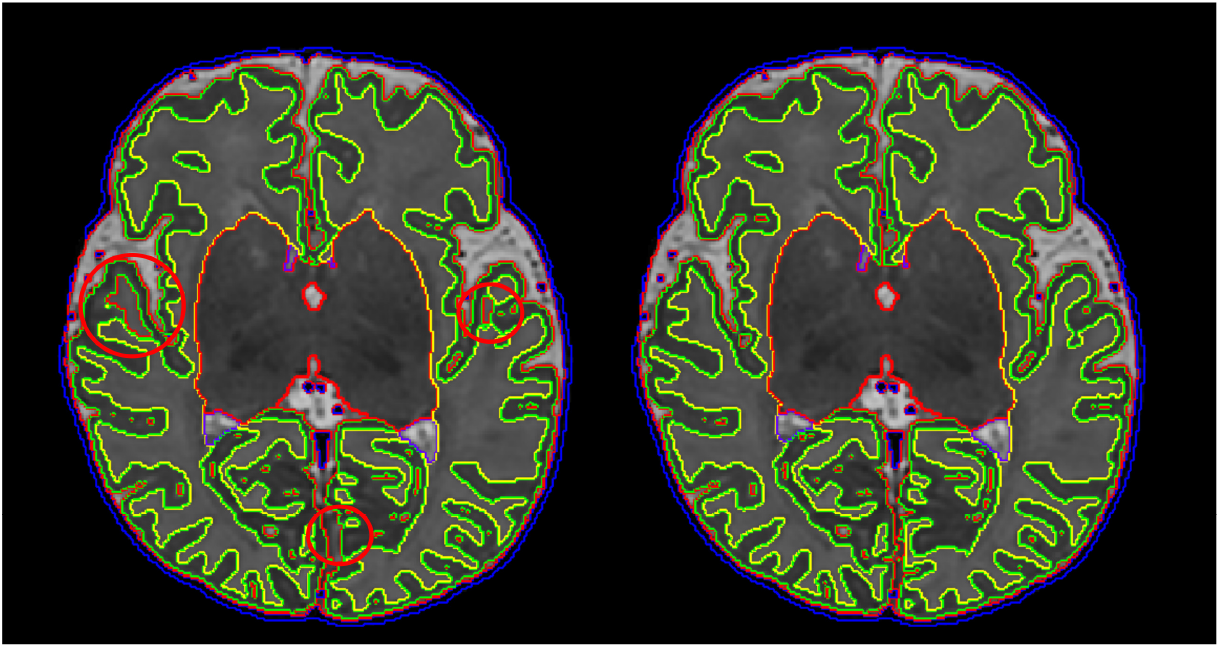
Segmentation of a subject T2 image with single-channel registration (left image) and multi-channel registration (right image). Affected areas are noted with a red circle.

**Figure 7:**
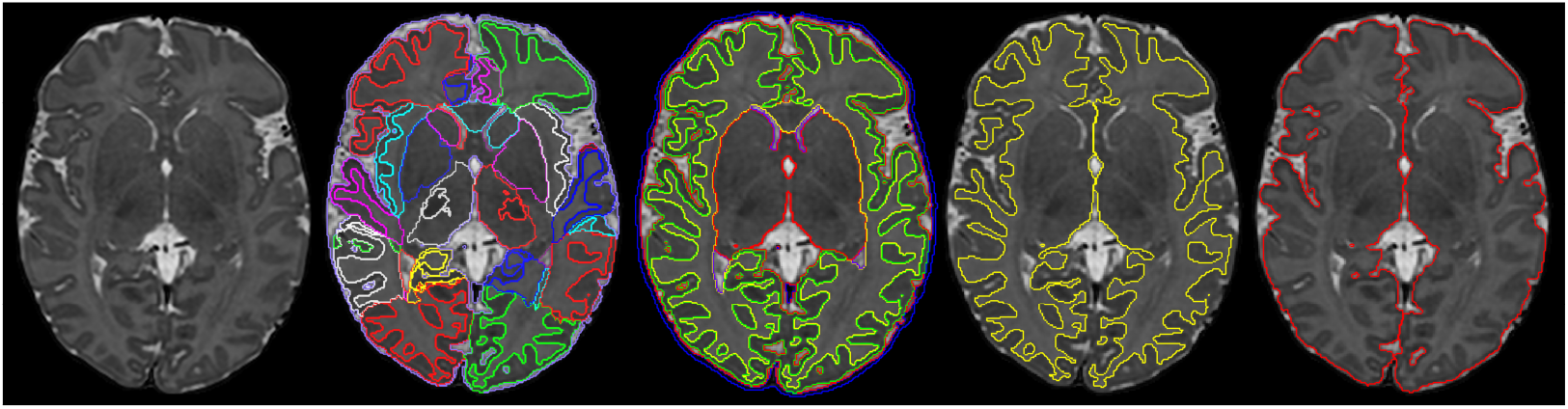
Example segmentation using Draw-EM. The subject T2, label segmentation, tissue segmentation, white surface and pial surface mask are presented from left to right.

**Figure 6:**
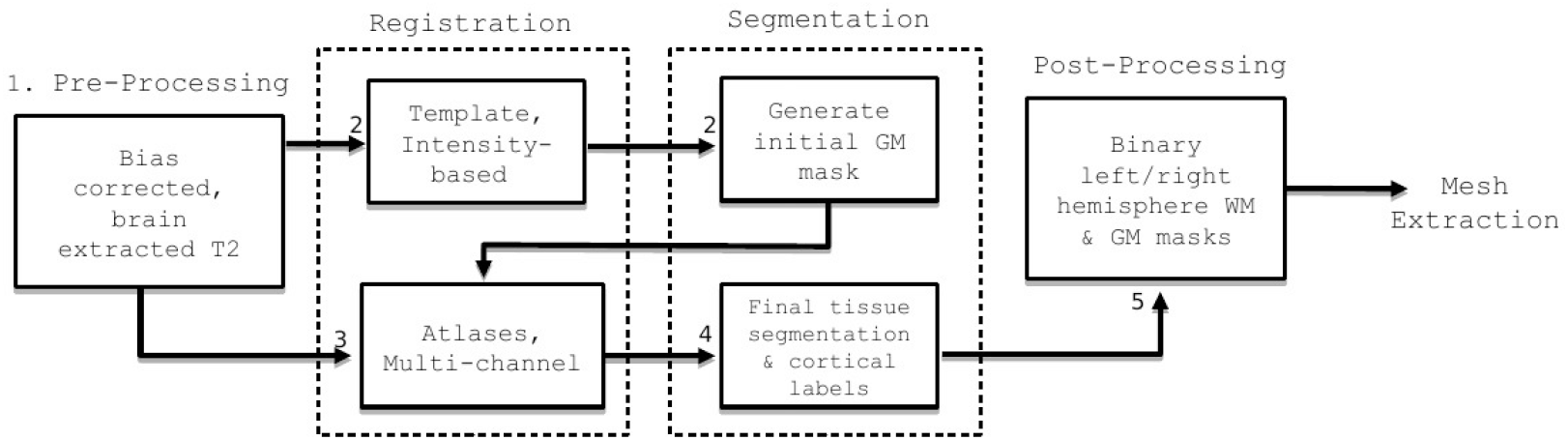
The major steps of the Draw-EM pipeline are: 1) Pre-processing. The original MRI is brain-extracted and corrected for intensity inhomogeneity; 2) The preterm atlas template is registered to the bias-corrected T2 image and an initial (tissue) segmentation is generated; 3) The GM probability map obtained from the initial segmentation is used together with the MR intensity image as different channels of a multi-channel registration of the labelled atlases by (Gousias et al., 2012) to the subject. 4) Regional labels are segmented based on the labels propagated from the atlases; 5) Segmented labels are merged in different granularities to further produce the following files: tissue labels, left/right white surface masks and left/right pial surface masks.

The labelled atlases (Gousias et al., 2012) are registered to the subject using a multi-channel registration approach, where the different channels of the registration are the original intensity T2-weighted images and GM probability maps. These GM probability maps are derived from an initial tissue segmentation, performed using tissue priors propagated through registration of a preterm probabilistic tissue atlas Serag et al. (2012) to each term-born subject. Preterm subjects typically have increased CSF volume over term neonates (Makropoulos et al., 2016), and this is reflected in the preterm template Serag et al. (2012). Therefore, to account for the effects of mis-registration resulting from the different amounts of CSF (see Fig. 4), we use a trimmed version of the preterm template (eroding CSF intensities from the template image) to perform the registration and initial tissue segmentation. Afterwards, these initial GM maps are then used along with the intensity T2-weighted images in the multi-channel registration of the labelled atlases (Gousias et al., 2012). Fig. 5 presents the benefit of using a multi-channel registration approach instead of a single-channel registration using only the intensity. It has been found that inclusion of the GM probability maps improves the accuracy of the alignment of the cortical ribbon, despite the large differences in cortical folding that are observed throughout the developing period. For more details on the individual parts of the original segmentation pipeline we refer the reader to Makropoulos et al. (2014).

### 3.3. White, Pial and Midthickness Surface Extraction

Following segmentation, white-matter mesh extraction is performed by fitting of a closed, genus-0, triangulated surface mesh (convex hull) onto the inferred white-matter segmentation boundary. However, the segmentation may include WM holes (WM misclassified as CSF) and undetected sulci (CSF misclassified as WM) due to partial volume effects. We therefore refine the shape of the mesh by incorporating intensity information from the bias-corrected T2 and (if available) T1 images (Schuh et al., 2017). This takes into account local geometry and favours sheet-like boundaries over thin bridges, or small pockets caused by CSF mislabelled as WM. Examples of the improvements gained by intensity based refinement, over a straightforward fit to the segmentation boundary, are shown in Fig. 8.

**Figure 8:**
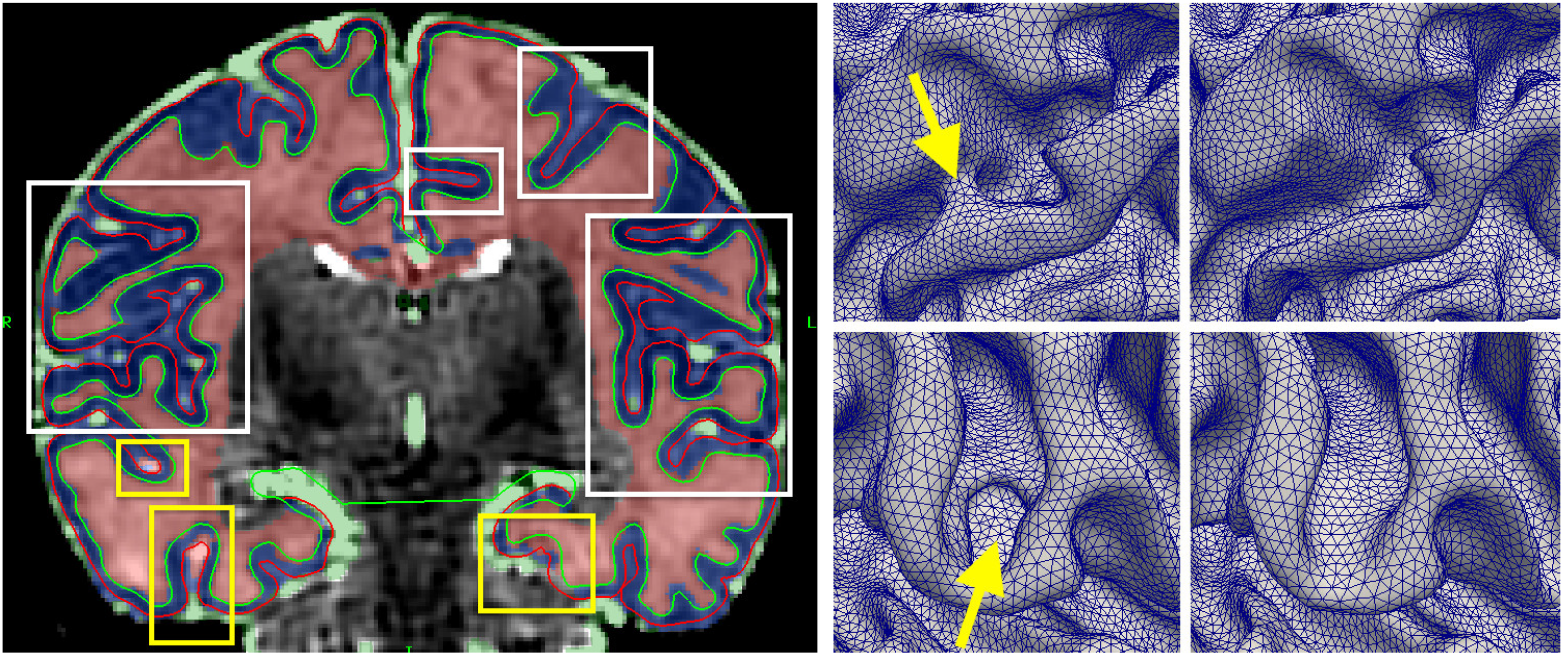
Segmented T2 image intersected by white and pial surfaces (left). White boxes outline corrections in areas where CSF appears dark due to partial volume effects, and yellow boxes corrections in areas where the CSF has been mislabelled as WM. Zoom of white surface mesh before (middle) and after (right) edge-based refinement. The top row demonstrates correction of a sulcus by moving the surface inwards, and the bottom row correction of a segmentation hole by moving the surface outwards.

In Schuh et al. (2017), the deformation of the initial genus-0 surface, inwards towards the white-grey tissue boundary, is governed through a trade-off between external and internal forces. Specifically, the external force seeks to minimise the distance between the placement of the surface mesh vertices and the locations of the tissue boundaries, and this is regularised by three different internal forces, which seek to: 1) enforce smoothness by reducing curvature; 2) flatten creases in the mesh along sulcal banks; and 3) discourage mesh intersections by introducing a repulsion force between adjacent nodes. In addition to the repulsion force, self-intersection is further prevented by predefining a minimum distance (0.1 mm) that the mesh triangles can reach between each other. If the new movement of a triangle moves it to a distance less than the minimum allowed distance to any other triangle in the mesh, this movement is prevented by limiting the total force of its vertices.

Surface reconstruction proceeds over two stages, where, in the first stage, external forces are derived from distances to the tissue segmentation mask, and in the second stage, boundaries are refined using external forces derived from intensity information: specifically, the force attracts each vertex node towards the closest WM/cGM edge in the normal direction.

A median filtering and Laplacian smoothing of surface distances is performed to reduce the influence of small irregularities in the segmentation boundary. Small holes in the segmentation manifest themselves in the surface distance map as small clusters of supposedly distant points as seen left of Fig. 10. These are filled in to avoid the surface mesh to deform into them.

The closest WM/cGM edge in the second stage is found by analysing the one-dimensional intensity profile and directional derivative sampled at equally-spaced ray-points. Examples are shown in Fig. 9. Here, a white-matter boundary edge occurs between a maximum with WM intensity, followed by a minimum with cGM intensity (left/right of green vertical line). Starting at the ray centre (red vertical line, yellow dot), a suitable edge is found by searching both inwards and outwards from the node until either a WM/cGM edge is found, a maximum search depth is exceeded, or the ray intersects the surface.

**Figure 9:**
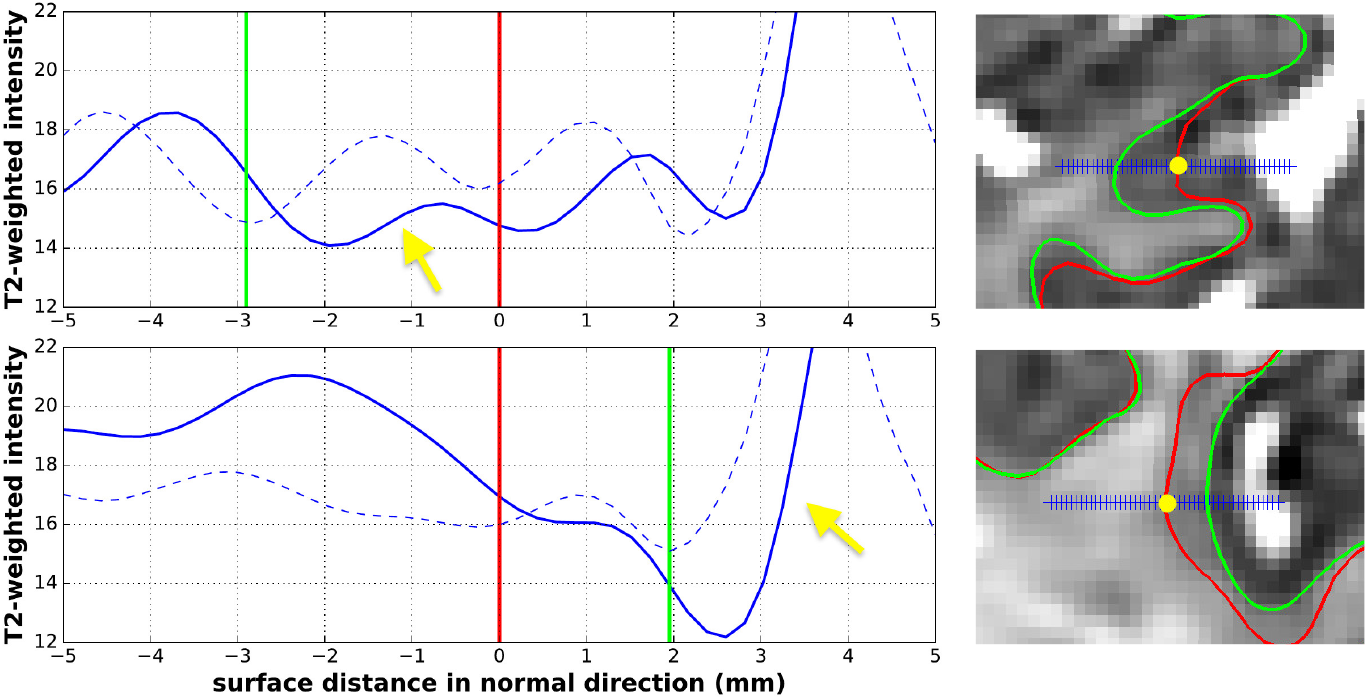
T2 signal intensity after bias correction (blue curve) and derivative (dashed curve) sampled at blue crosses. The yellow dot marks the mesh node on the initial surface (red). The green contour depicts the white-matter surface. Yellow arrows point at the cGM/CSF edge of the pial surface.

Next, the pial surface is obtained by deforming the white-matter mesh towards the cGM/CSF boundary using the same model but modifying the external force to search for the closest cGM/CSF image edge outside the white-matter mesh. These edges correspond to a positive derivative in normal direction of the T2 intensity (yellow arrows in Fig. 9). When no such edge is found, e.g. within a narrow sulcus due to partial volume, the opposing sulcal banks expand towards each other until stopped by the non-self-intersection constraint. Finally, a midthickness surface is generated half-way in-between by averaging the positions of corresponding nodes. Please refer to Schuh et al. (2017) for more details on the surface reconstruction.

### 3.4. Surface Inflation and Spherical Projection

In a separate process, the white matter surface is inflated using a reimplementation of the inflatable surface model of FreeSurfer (Fischl et al., 1999a). This uses an inflation energy functional that consists of an inflation term, that forces the surface outwards (for vertex points in sulcal fundi) and inwards (for points on gyral crowns), and a metric-preservation term that preserves distances between neighbouring points. Surfaces are expanded until they fulfil a pre-set smoothness criterion (Fischl et al., 1999a).

As a by-product of the surface inflation, a measure of average concavity/convexity is obtained for each vertex. This represents the change of a node’s position in normal direction, and is proportional to the depth of major sulci.

**Figure 10:**
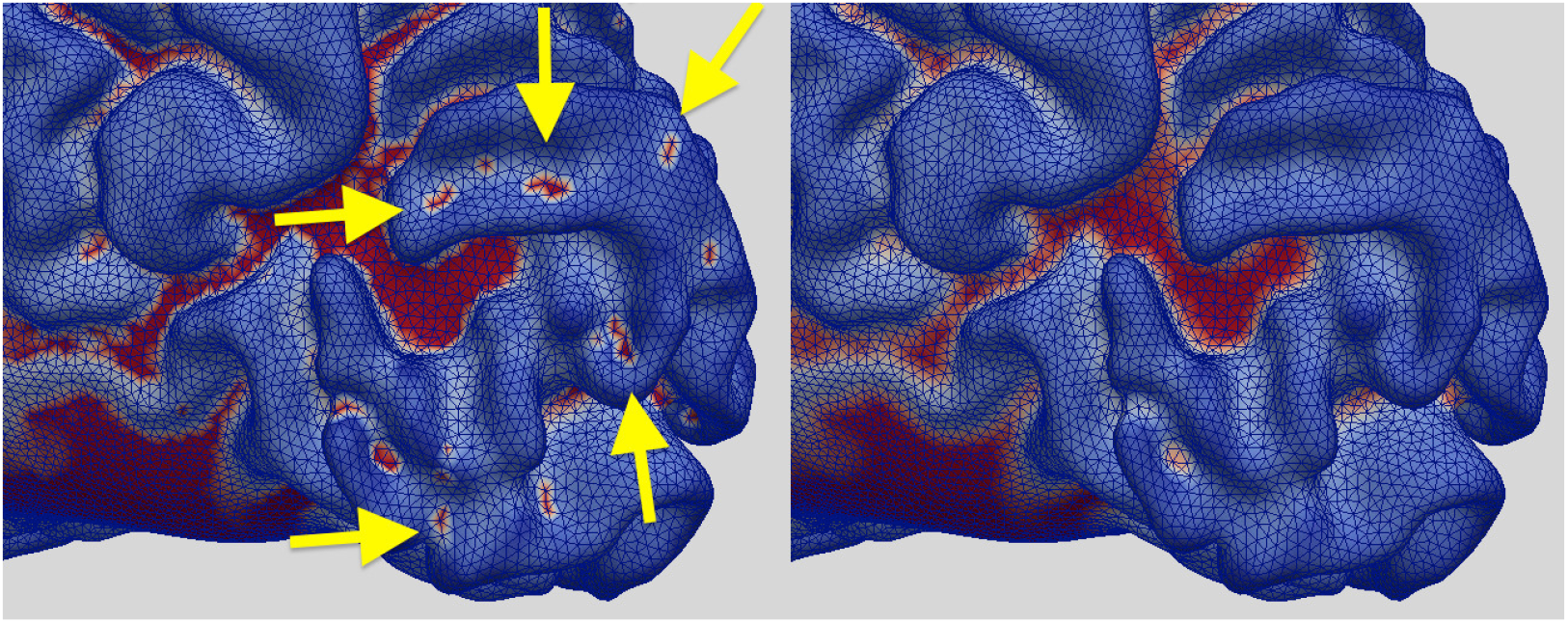
WM segmentation boundary distance in normal direction, before clustering based hole filling (left), and after the hole filling and smoothing (right). A number of holes in the segmentation are indicated by yellow arrows.

In a final step, inflated meshes are projected onto a sphere using spherical Multi-Dimensional Scaling (sMDS) as proposed by (Elad et al., 2005). This computes an embedding, or projection, of each inflated mesh onto a sphere, whilst minimising distortion between mesh edges. More specifically, the projected positions of vertex points on the sphere are optimised so as to preserve the relative distances between points, as measured in the initial inflated mesh space. In each case, the distances (measured for the sphere and the input mesh) are estimated as geodesics, and are computed using a fast marching method for triangulated domains (Kimmel and Sethian, 1998). Note, the input mesh must first be scaled so that its surface area is equal to that of the unit sphere.

The benefit of sMDS is that the optimisation of the spherical projection is performed directly on spherical coordinates, while in FreeSurfer the distances are optimized in 3D. To evaluate the performance of sMDS relative to FreeSurfer we compared areal and edge distortion distributions for vertex spacing in the original white matter mesh relative to the sphere. Here, areal distortions measure change in mesh faces area estimated as 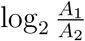 (for triangle *A*_1_ in white mesh, and equivalent triangle *A*_2_ on the sphere). Edge distortions reflect change in relative vertex spacing and are estimated as 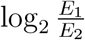 (for edge length *E*_1_ in white mesh, and equivalent *A*_2_ on the sphere). Distortion distributions are averaged across all subjects and both hemispheres, and results are shown in Fig 11. Results show that while distortions for the sMDS method slightly exceed that of FreeSurfer, they are in a comparable range.

**Figure 11:**
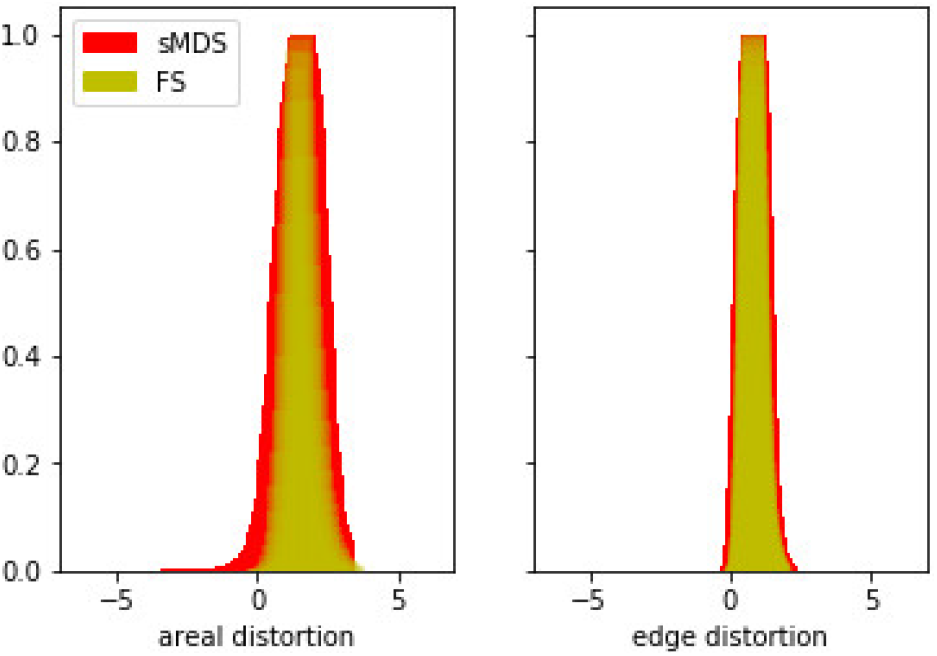
Spherical projection distortions for FreeSurfer (FS) and spherical MDS (sMDS)

### 3.5. T1-T2 registration

As the neonatal pipeline is based on T2 image processing rather than T1 (as for the adults), each T1 image was rigidly registered to its T2 image pair. Similarly to the HCP, registration is performed with Boundary-Based Registration (BBR) Greve and Fischl (2009) to estimate the 6 degrees of freedom (DOF) rigid registration parameters. However, directly computing the registration with BBR resulted in misalignments in 16.8% of the scans (69 out of the 411 cases with both T1/T2 scans). Due to these mis-alignment issues, we performed an initial rigid registration of the T1 to an eroded T2 excluding the cGM and CSF (including the WM and sub-cortical structures). BBR is then initialised with the parameters from this initial registration, which provides further minor improvements. The combined rigid registration, initial rigid followed by BBR, resulted in only one case with minor residual misalignment to its corresponding T2 image (0.2% of the scans).

### 3.6. Feature Extraction

The processes of surface extraction and inflation generate a number of well known feature descriptors for the geometry of the cortical surface. These include: surface curvature estimated from the mean curvature (or average of the principal curvatures) of the white matter surface; cortical thickness estimated as the average distance between a) the Euclidean distance from the white surface to the closest vertex in the pial surface and b) the Euclidean distance from the pial surface to the closest vertex in the white surface; and sulcal depth which represents average convexity or concavity of cortical surface points estimated during the inflation process.

In addition to geometric features, the pipeline also estimates correlates of cortical myelination using the protocol described in Glasser and Van Essen (2011); adopted by the HCP. These feature maps are estimated from the ratio of T1-weighted and T2-weighted images and represent an architectonic marker that reliably traces areal boundaries of cortical areas and correlates with cytoarchitecture and functionally distinct areas in the cortex Glasser and Van Essen (2011). As the neonatal pipeline is based on T2 image processing rather than T1 (as for the adults), neonatal T1/T2 myelin maps are estimated by first rigidly registering each T1 image to its T2 image pair. The ratio is estimated from the original T2 image and the transformed T1 image (prior to bias correction). Following this, the T1/T2 ratio is projected onto the midthickness surface, using volume-to-surface mapping (Glasser et al., 2013).

Further, regional labels (from the method described in Gousias et al. (2012)) are automatically generated during segmentation. These regional labels are further projected onto the cortical surface. The closest voxel in the volume with respect to each vertex in the midthickness surface is used to propagate the label to that vertex. The projected labels are consequently post-processed with filling of small holes and removal of small connected components.

Fig 12 depicts exemplar surfaces for three subjects aged 32, 36 and 40 weeks PMA (at time of scan) together with cortical labels, sulcal depth, cortical thickness, mean curvature, and T1/T2 myelin maps. This demonstrates the considerable differences in surface geometry and myelination over the developmental period covered by this cohort.

**Figure 12:**
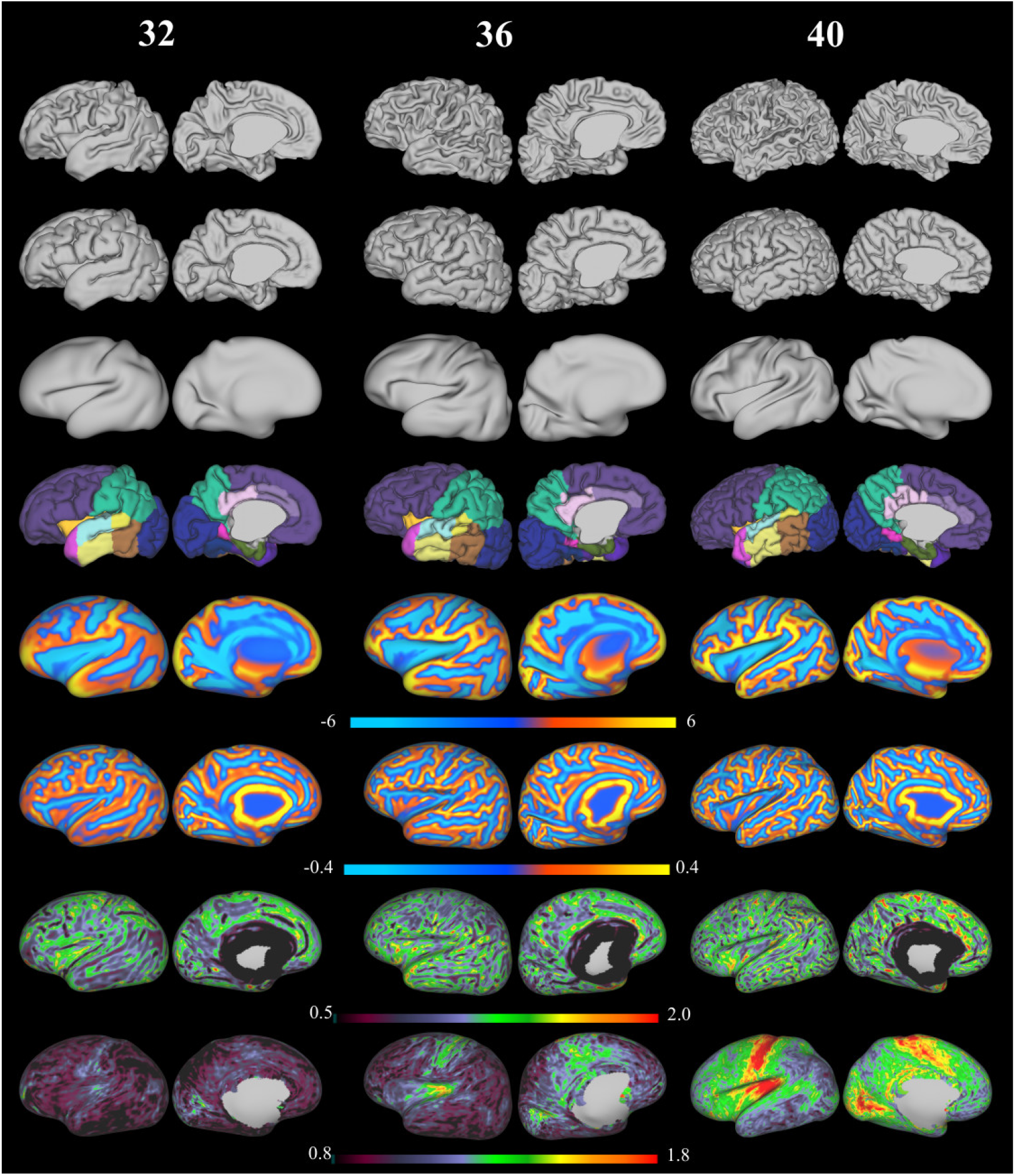
Exemplar surfaces and feature sets for three individuals scanned at 32, 36 and 40 weeks PMA (left hemisphere). From top to bottom: white surface, pial surface, inflated surface, cortical labels, sulcal depth maps, mean curvature, cortical thickness and T1/T2 myelin maps (all features shown on the very inflated surface)

## 4. Quality Control (QC)

The quality of the pipeline was assessed by manually scoring a sub-set of randomly selected images from the cohort. We performed three independent scorings for the different parts of the pipeline: image reconstruction, segmentation and surface reconstruction. 160 images were used for the image reconstruction and segmentation QC. A sub-set of 43 images was then used for the surface reconstruction QC due to the more manually intensive and time consuming scoring of cortical details. All images and surfaces were scored independently by two expert raters (one neuro-anatomist, one methods specialist) to assess inter-rater agreement. The image and segmentation QC was performed by AM and SJC and surface QC by AM and JS. The results are presented in the following sections.

### 4.1. Image QC

Quality checking of neonatal images was performed by first binning all subjects’ PMA (at scan) into weeks (in the range: 37–44 weeks) and selecting 20 images at random from each weekly interval. This yielded a total of 160 images. Note, the relatively low limit of 20 per week was selected due to the limited amount of images (57 out of the total 492 scans, less than 20 per age) acquired at earlier scan ages (29 – 36 weeks). The protocol for manual scoring of the image quality is presented in Fig. 13. Scoring was performed by visual inspection of the whole 3D volume slice by slice. Poor quality images were rated with score 1. Images with significant motion were rated with score 2. Images with negligible motion visible only in a few slices were rated with score 3. Images of good quality with no visible artefacts were rated with score 4.

**Figure 13:**
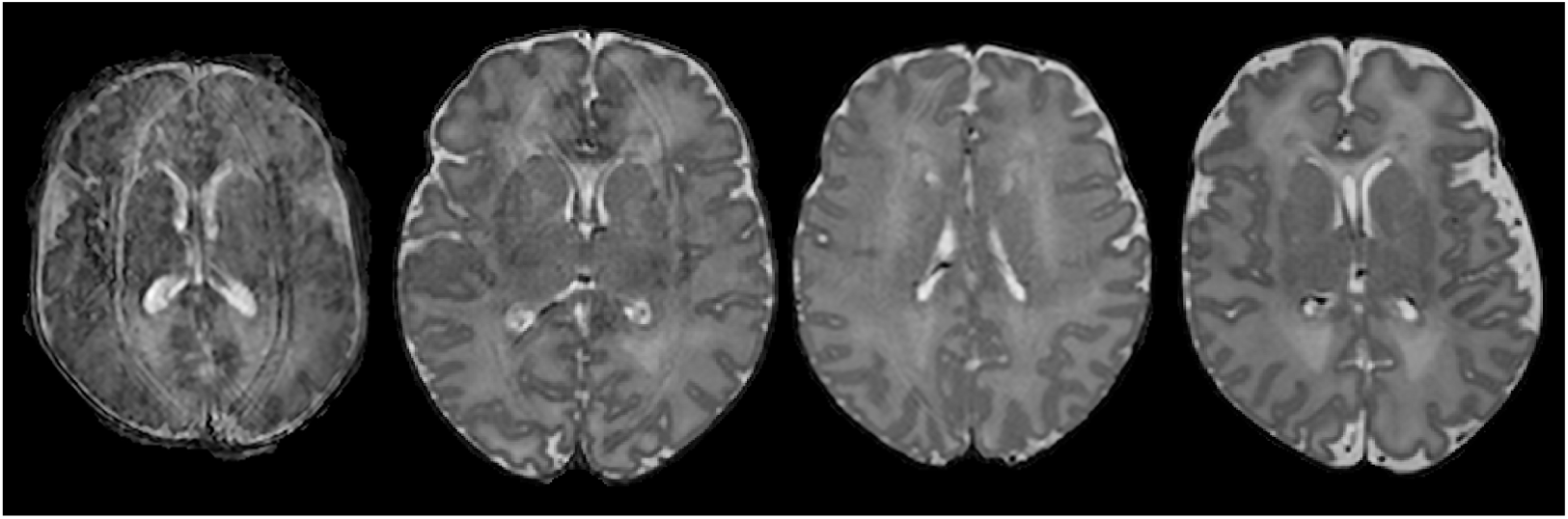
Manual QC score of image quality. From left to right: poor image quality - score 1, significant motion - score 2, negligible motion - score 3, good quality image - score 4.

Fig. 14 presents the percentage of images rated with the different scores by the two reviewers. On average, 90% of the images were considered to have negligible or no motion (score 3 and 4) and only 2% of the images had to certainly be discarded due to poor image quality according to the raters. The raters’ scoring was the same in 78% of the images and within a difference of one scoring point in 99% of the images.

**Figure 14:**
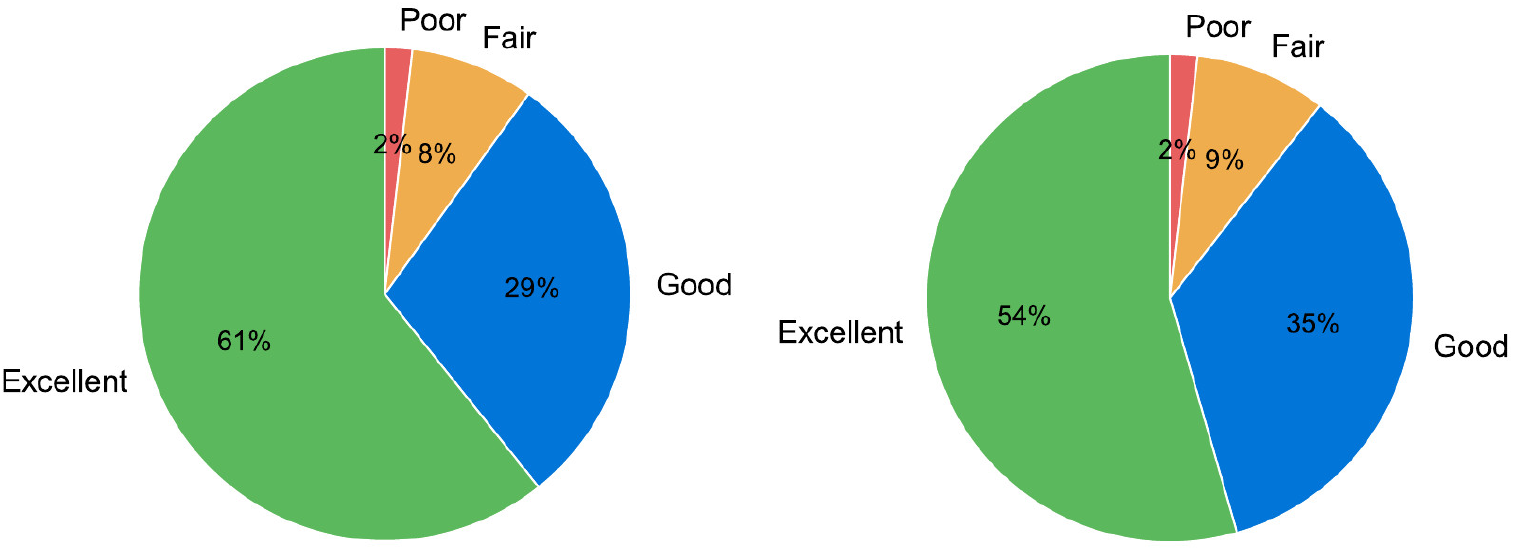
Image scores from the 2 raters (based on 160 images).

### 4.2. Segmentation QC

Following image QC, tissue segmentation masks were evaluated for the same set of subjects. Quality was assessed using the protocol presented in Fig. 15. Specifically, images that were poorly segmented were rated with score 1; images with regional errors were rated with score 2; images with localised segmentation errors were rated with score 3; images with segmentations of good quality with no visible errors were rated with score 4.

**Figure 15:**
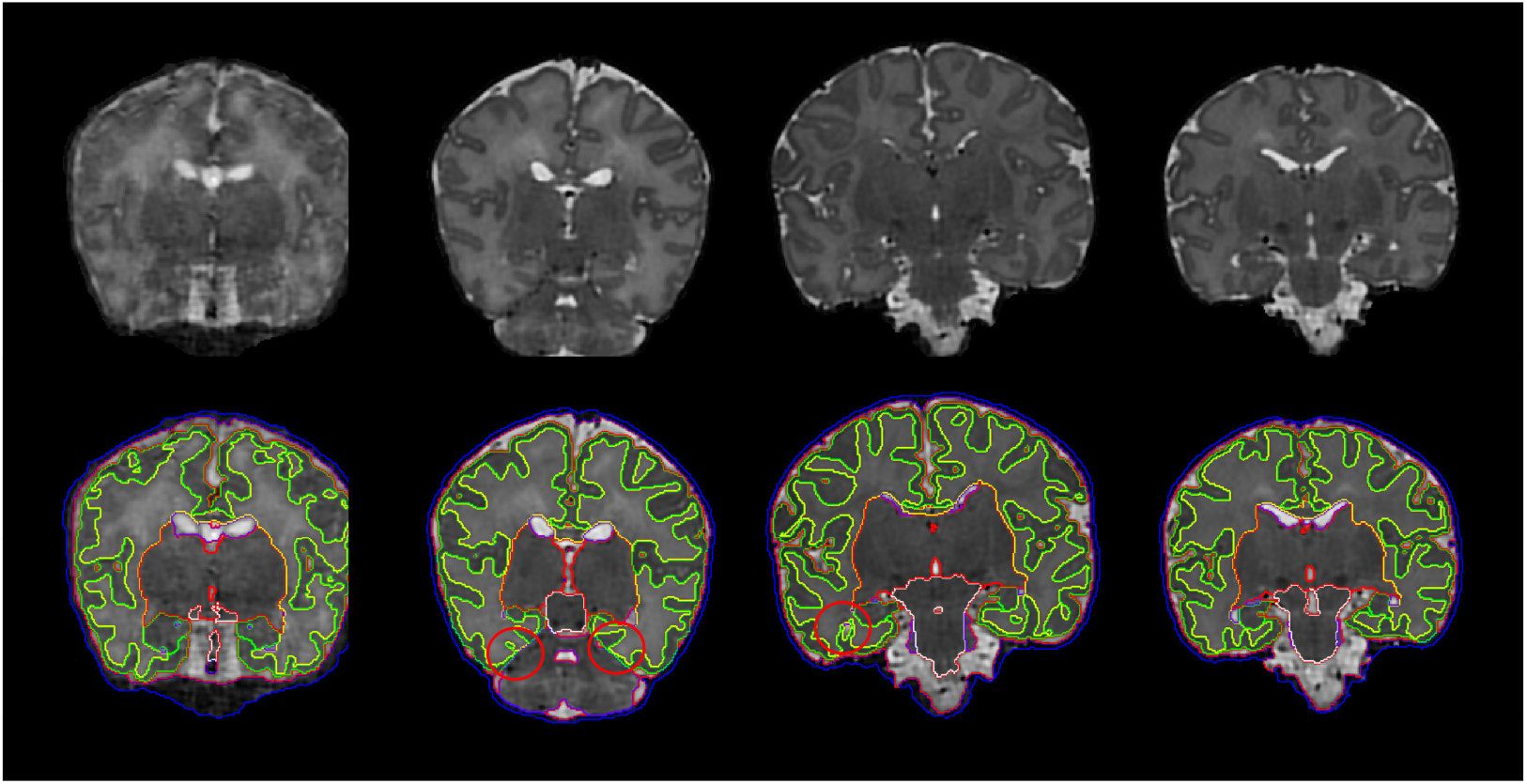
Manual QC score of segmentation quality. From left to right: poor segmentation quality - score 1, regional errors - score 2, localised errors - score 3, good quality segmentation - score 4.

Regional errors often occurred at the interface between the cerebellum and the inferior part of the occipital and temporal lobe, where part of the cortical ribbon was mislabelled as cerebellum due to poor image contrast between the regions; another source of regional problems was when WM was mislabelled as cGM or CSF due to partial volume effects. By contrast, localised errors typically occurred when the CSF inside a sulcus was mislabelled as WM due to partial voluming effects.We further refer the reader to Makropoulos et al. (2014) for additional validation of the segmentation method.

The scores attributed by the two raters are presented in Fig. 16. Poor segmentation occurred only in 5 scans (3% of cases) and these were related to poor image quality or significant motion (image score 1 and 2). Regional (24% of cases) and localised errors (64% of cases) were present in the majority of the cases. The scoring of the raters agreed in 71% of the images and was within a difference of one scoring point in 99% of the images.

**Figure 16:**
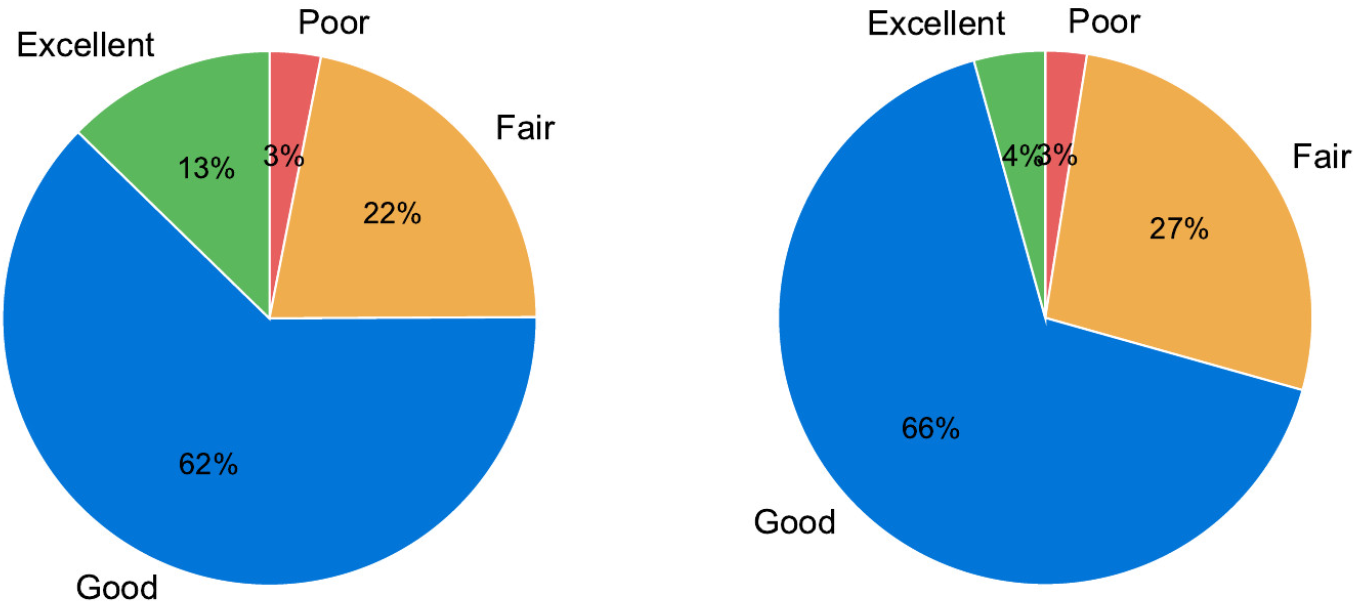
Segmentation scores from the 2 raters (based on 160 images).

Furthermore, we assessed the improvements with the proposed segmentation. We manually inspected segmentations of all the 492 scans obtained with: a) the proposed segmentation method and b) the tissue segmentation method outlined in Makropoulos et al. (2012), that uses the original template from Serag et al. (2012) for the atlas priors, as baseline (Makropoulos et al. (2012) presented the most accurate results in the NeoBrainS12 challenge (Išgum et al., 2015)). Fig. 17 presents the occurrence of the different segmentation problems presented in Section 3.2 with the baseline and the proposed method: top part of cGM missing (see Fig. 4), CSF inside WM (see Fig. 5), hyper-intense WM misclassified as ventricles (see Fig. 3), hypo-intense WM misclassified as cGM (see Fig. 3). The segmentation method with the proposed modifications has diminished the occurrences of this problems (top part of cGM missing in 1.6% compared to the baseline 23.8% of cases, CSF inside WM in 1.4% compared to 36.4%, hyper-intense WM misclassified as ventricles in 7.7% compared to 90.7% and hypo-intense WM misclassified as cGM in 0% compared to 84.6%). These problems, where still present in the proposed method, are typically reduced in extent.

**Figure 17:**
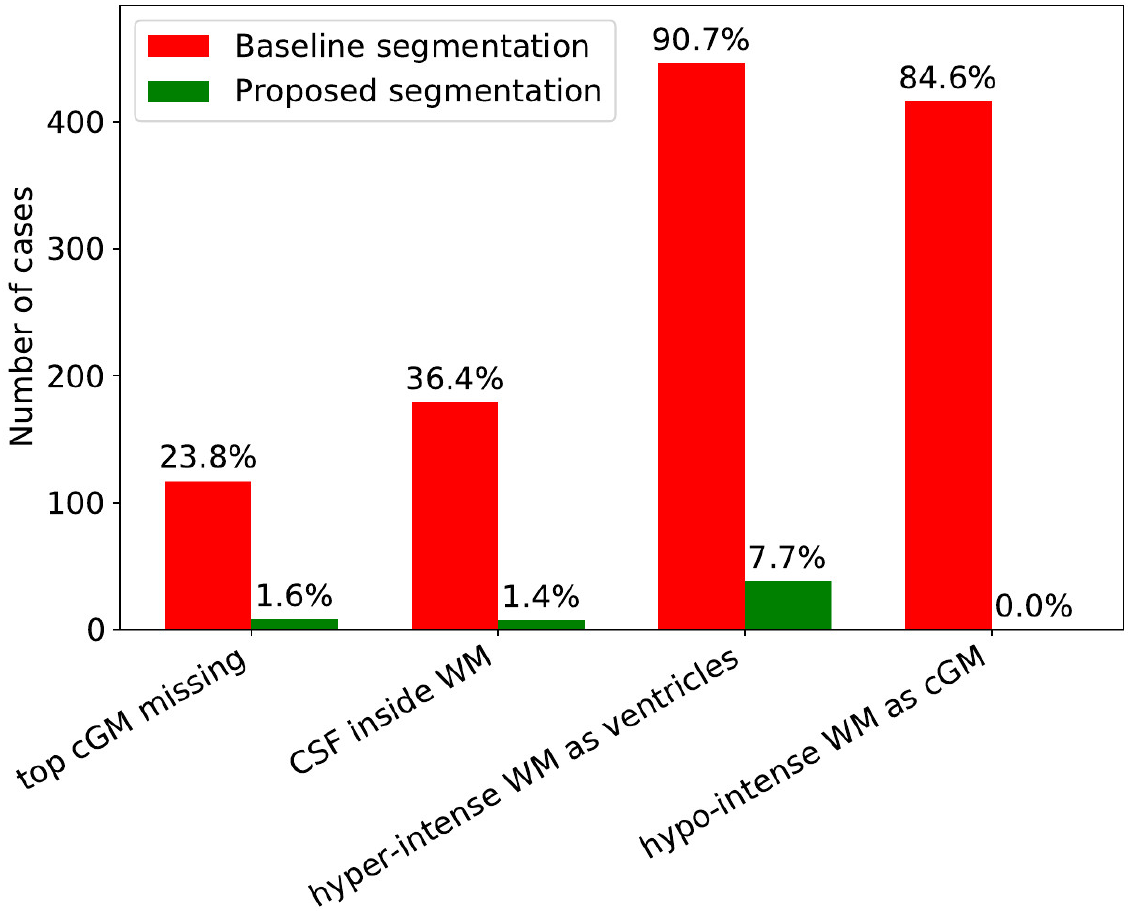
Occurrence of segmentation problems with the baseline segmentation method (Makropoulos et al., 2012) and the proposed segmentation (based on 492 images).

The tissue segmentations presented additional errors that were not examined in the segmentation QC. These can be seen in Fig. 18. Narrow WM folds present in the medial temporal and occipital lobe, and gyri at the most superior part of the brain can be misclassified as cGM. This is due to partial volume effects and low local intensity contrast that can not be easily resolved by intensity alone without prior knowledge of the cortical morphology.

**Figure 18:**
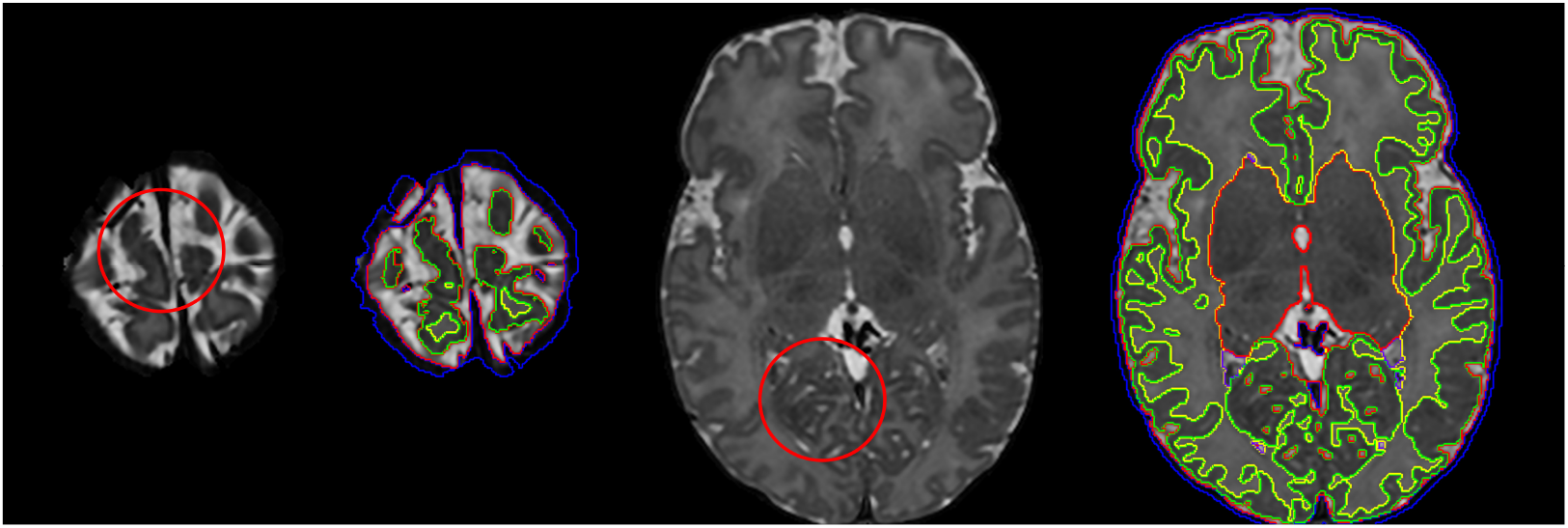
Additional errors remaining at the segmentations due to low intensity contrast and partial volume effects. WM at the superior part of the brain (left images) and narrow cortical folds at the medial part of the occipital lobe (right images) are occasionally misclassified as cGM.

Figures A.1, A.2 present example segmentations from the subjects included in this study.

### 4.3. Surface QC

Cortical surface quality was assessed by visual scoring of the reconstructed white surface, as this is the primary surface used in downstream functional and diffusion MRI processing. Visual surface QC by an expert was performed for a number of regions of interest (ROIs), automatically selected per subject. In order to specifically focus the surface QC on the improvements of the reconstructed surface over the tissue segmentation boundary produced from Draw-EM, we aimed to identify regions that deviate from this boundary. We additionally reconstructed each surface with an alternative surface extraction technique that faithfully follows the tissue mask segmentation boundaries (Wright et al., 2015). Surface QC then proceeded for ROIs where the surfaces extracted from the two techniques (Schuh et al. (2017) and Wright et al. (2015)) deviated most from each other. ROIs were viewed in image volume space as 3D patches (of size 50 × 50 × 50 mm) located at the centre of each cluster, with the white surface shown as a contour map (see Fig. 19 for examples of the ROI visualization). Due to the large number of patches, a subset of 43 images was used from the 160 images included for the image and segmentation QC. 20 ROIs were selected for each subject and were rated independently. Scoring of the surfaces was done by assigning a score from 1 to 4 for each ROI according to the protocol in Fig. 19: score 1 was rated as poor quality (this happened typically in cases where the contour substantially deviated from the cortical boundary or there were extensive missing gyri); score 2 was assigned in cases where the contour was close to the cortical boundary but there were obvious mistakes; score 3 was used for contours that were accurate but contained some minor mistakes; score 4 indicated an accurate contour tracing of the cortical boundary.

**Figure 19:**
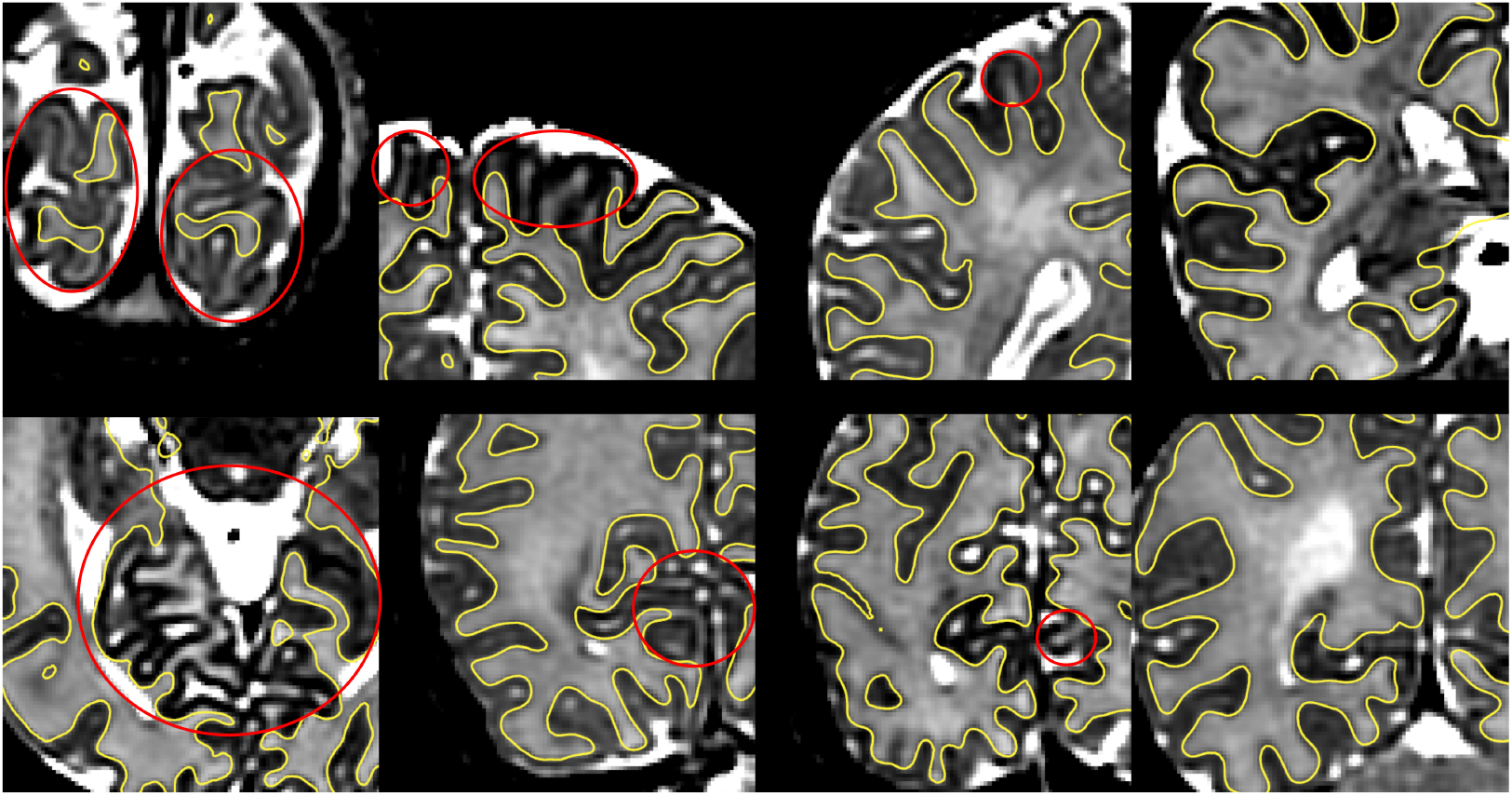
Manual QC score of surface quality. From left to right: poor surface quality - score 1, contour close to the cortical boundary but with obvious mistakes - score 2, minor mistakes - score 3, good quality surface - score 4.

Fig. 20 presents the results of the surface QC. The proposed method produced surfaces that were rated as accurate (score 3 or 4) in 85% of the ROIs with 49% presenting no visual mistakes, averaged between the two raters. The remaining percentage of ROIs (15%) were rated as having obvious mistakes but being close to the cortical boundary (score 1 or 2), with only less than 2% having poor quality (score=1). The scoring of the raters was the same in 52% of the ROIs and was within a difference of one scoring point in 93% of the ROIs. Fig. 21 presents the scoring for the different regions of the brain. With the exception of the temporal lobe, it can be observed that the raters score similar number of ROIs with score 1 and 2, but differ on the assignment of scores 3 and 4. On average, the occipital lobe presents the largest proportion of scores 1 and 2 (22%), followed by the temporal lobe (15%), the parietal lobe (11%), the frontal lobe (7%), corpus callosum (5%), insula (3%) and the cingulate gyrus (0%). Parts of the occipital and temporal lobe that contain narrow folds are hard to segment due to partial volume effects and low contrast, and errors in these regions are also propagated to the surfaces. Note, that in extreme cases of partial volume, tissue boundaries will also not be apparent to manual observers. In such cases, evaluators must make subjective choices that may explain the disagreement between raters.

**Figure 20:**
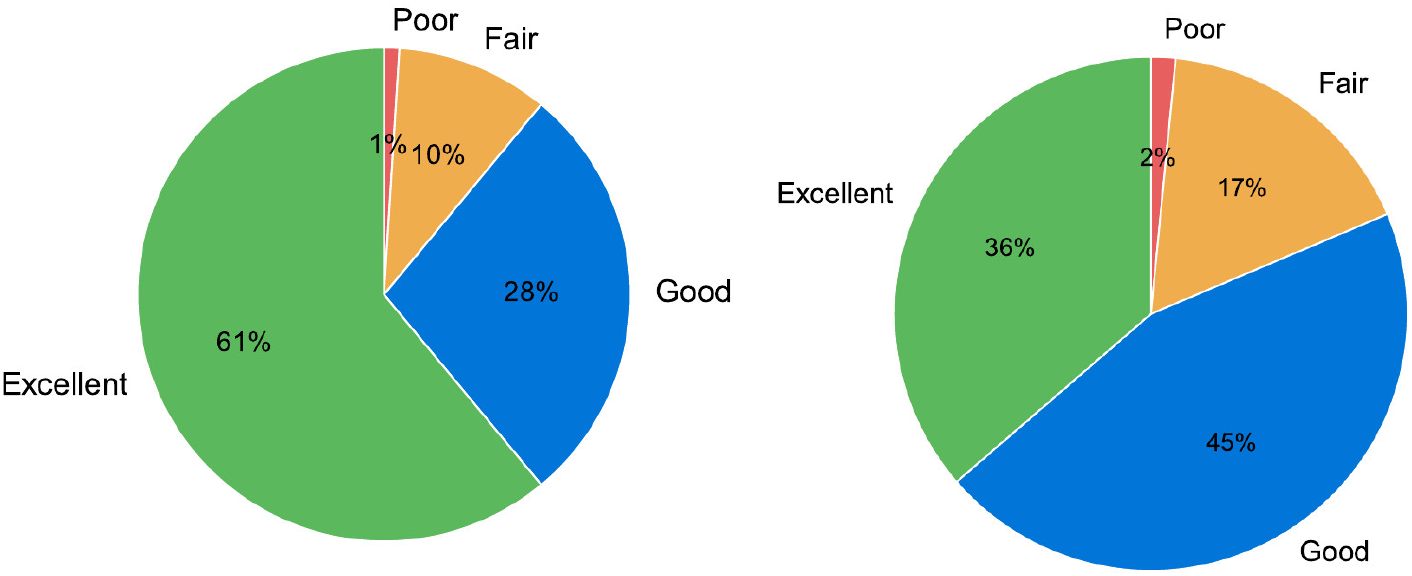
Scoring of surface ROIs from the 2 raters (based on 43 images with 20 ROIs per image).

**Figure 21:**
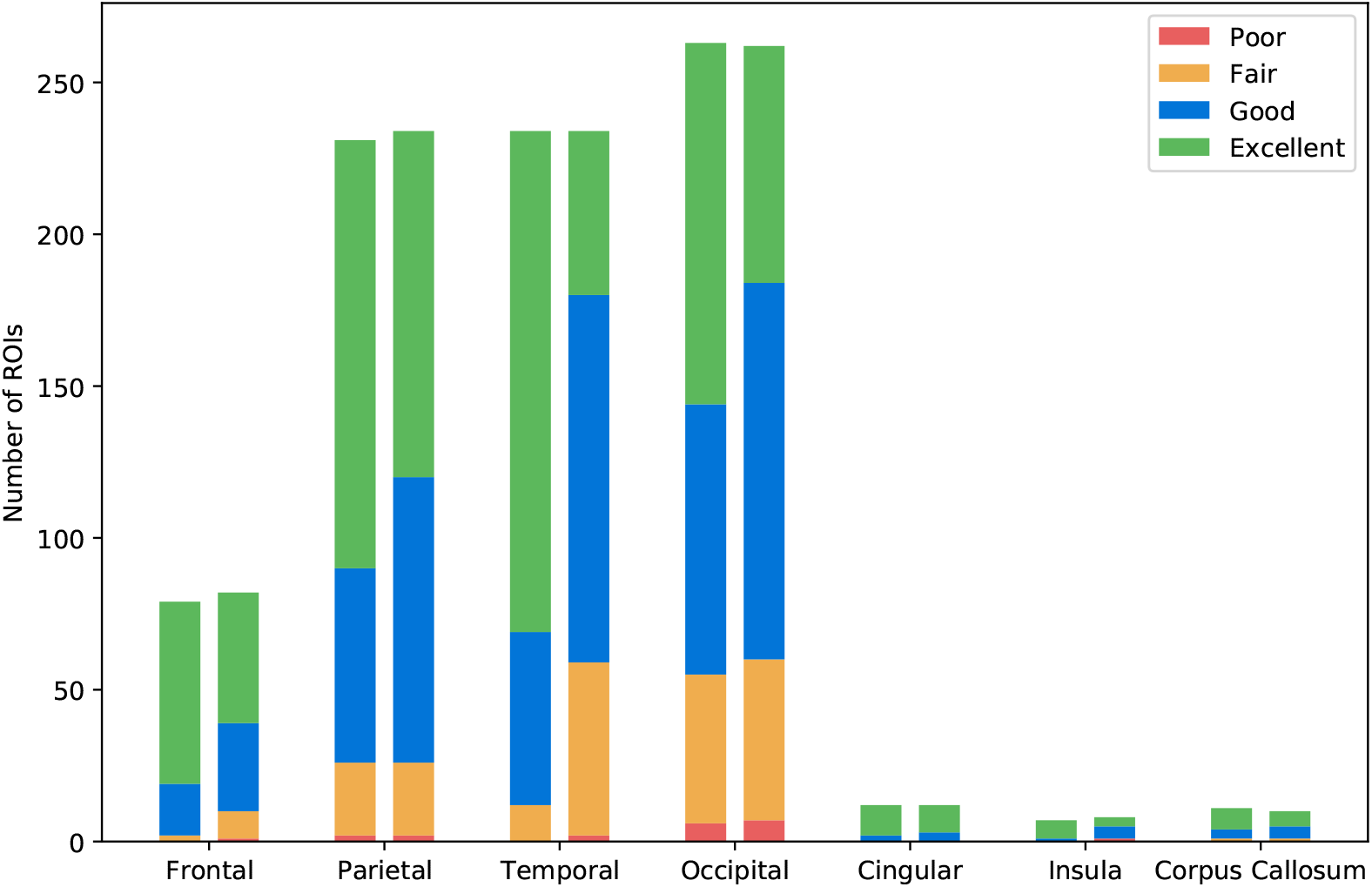
Regional surface scores from the 2 raters (based on 43 images with 20 ROIs per image).

Fig. 22 demonstrates examples of improvements with the proposed method over a surface extracted to faithfully follow the segmentation boundary. Figures A.1, A.4 and A.5 present example surfaces reconstructed from the data. Figures 23, 24 further present the regional and total cortical thickness of all the subjects, and average thickness maps across different ages at scan. Average thickness maps were calculated by registering the sulcal depth maps of the subjects with MSM (Robinson et al., 2014) to an average template constructed using the method described in Bozek et al. (2016b). Cortical thickness across the whole brain has a mean value of 1.1 mm, which increases across the age at scan. Cortical thickness measurements in the neonatal population have been previously reported in the literature in (Xue et al., 2007; Moeskops et al., 2013; Li et al., 2015a; Moeskops et al., 2015; Makropoulos et al., 2016; Geng et al., 2017) with different ranges between 1-2 mm. Differences between reported values can be due to different image acquisition, image segmentation and surface reconstruction methods. Moeskops et al. (2013), in contrast to other studies, present thickness values estimated based on manually segmented cortices. The estimated thickness in their study varies between 0.95–1.2 mm which is very similar to the obtained measurements here. Different parts of the neonatal brain, such as the major lobes (frontal, parietal, temporal, occipital), present differences in the thickness values Moeskops et al. (2013, 2015); Geng et al. (2017). Li et al. (2015a) present thickness values at 0,1 and 2 years of age and report higher values in the frontal and temporal lobe, and lower values in the parietal and occipital lobe. A similar trend can be observe here (temporal lobe: 1.15 mm, frontal lobe: 1.13 mm, parietal: 1.1 mm, occipital: 1.05 mm).

**Figure 22:**
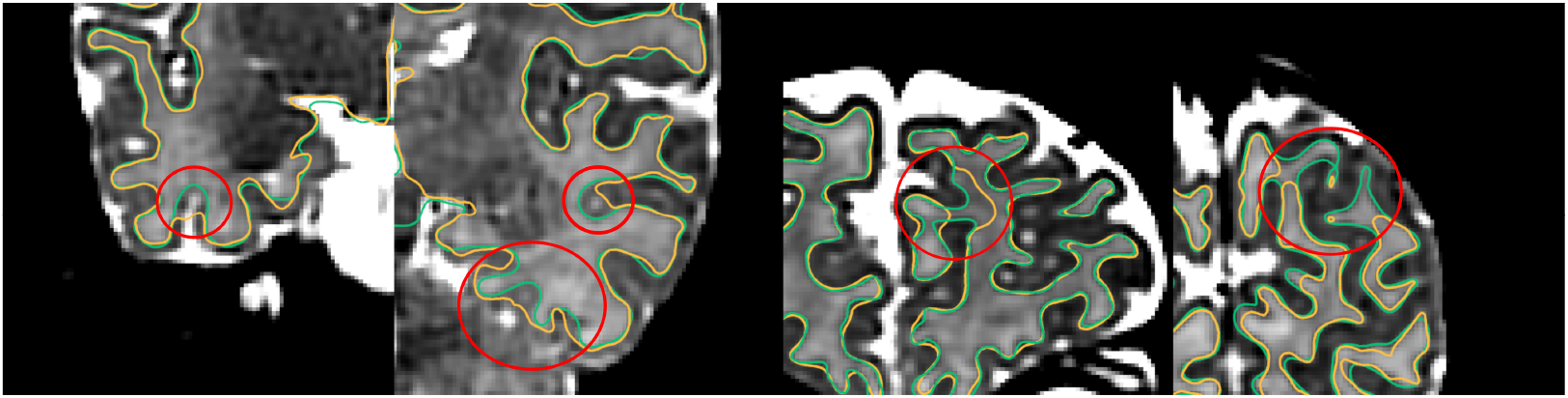
Comparison of surfaces produced with the proposed method (overlaid with green colour) over surface extraction that follows the tissue segmentation boundary produced from Draw-EM (overlaid with yellow colour).

**Figure 23:**
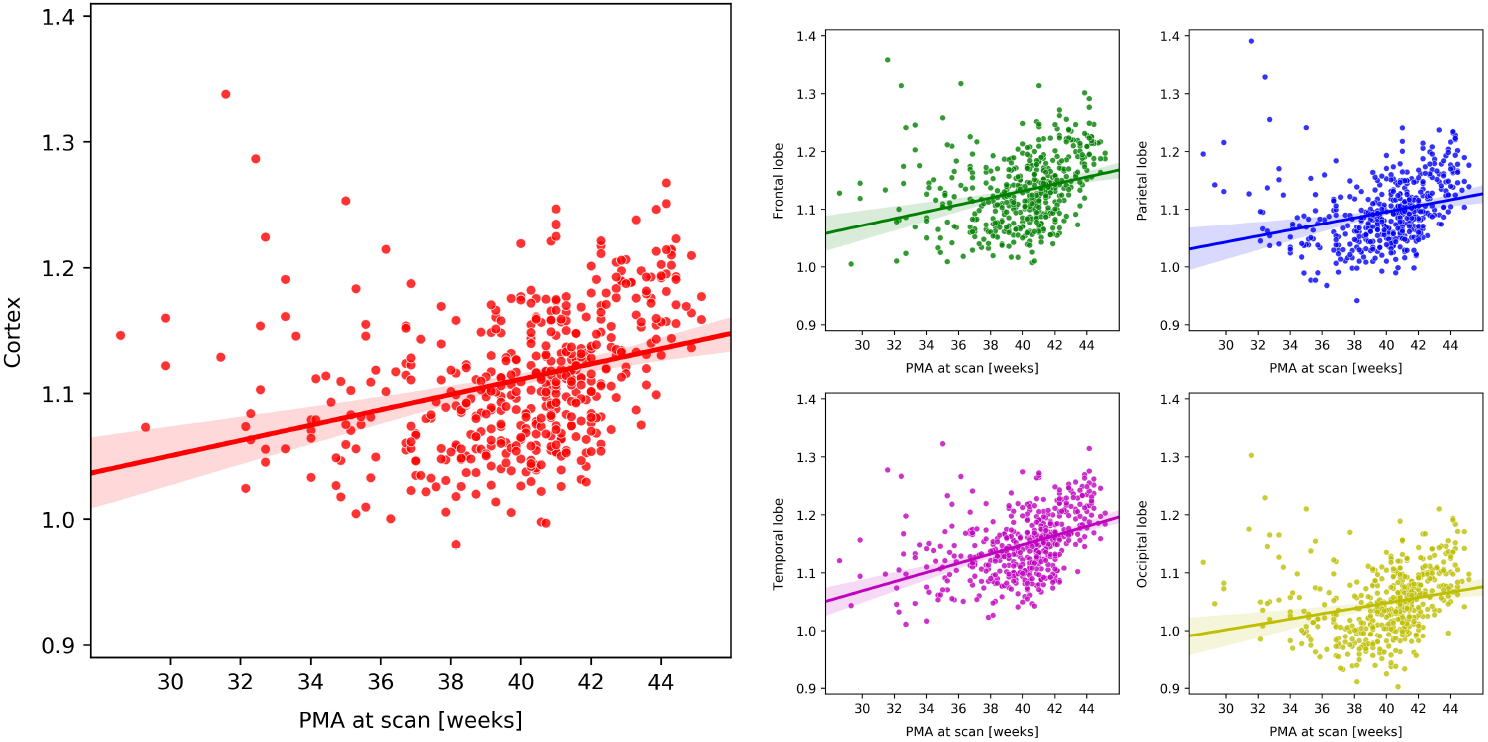
Average thickness (mm) of the cortex and different lobes across different ages at scan.

**Figure 24:**
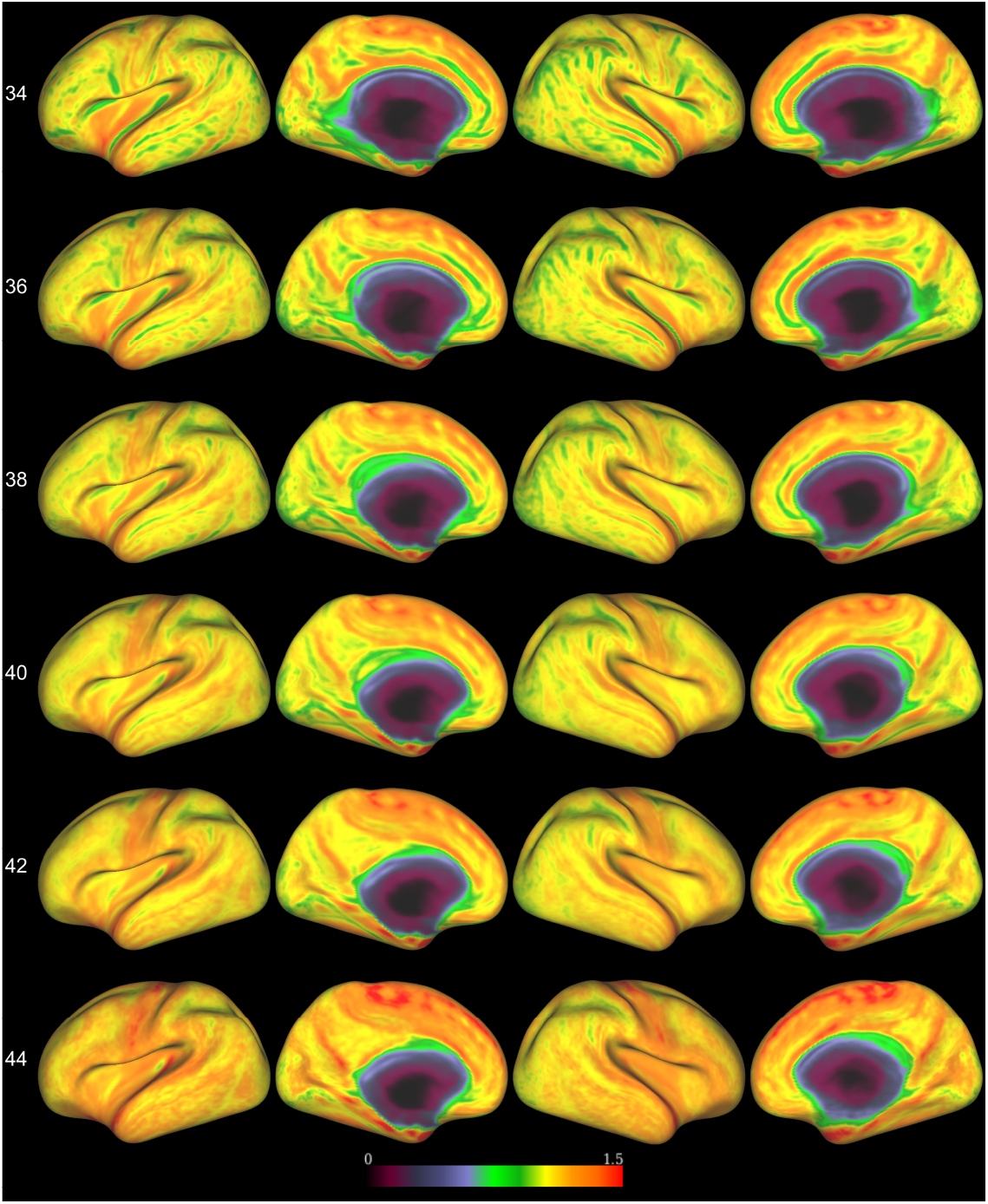
Average thickness maps of the left and right hemisphere across different ages at scan.

It should be noted that the adopted surface reconstruction model entails a number of parameters (e.g. weight of external force, strength of smoothness constraints imposed by internal forces and minimum vertex distance). Due to lack of ground truth these parameters were set based on visual inspection of the surface renders in 3D, and super-imposing them on the image. These parameters are likely specific to the dHCP cohort, and reflect subjective choices with respect to the balance of smoothness, relative to goodness of fit of the boundaries. We recommend users of the pipeline re-optimise parameters for new cohorts.

## 5. Comparison to HCP Pipelines

The dHCP pipeline has been inspired by the HCP (Glasser et al., 2013). However, there are several key differences between the pipelines, as highlighted in Table 3. Neonatal subjects are imaged during natural sleep. Therefore, total scanning time for all structural, diffusion and rfMRI scans is markedly reduced relative to the HCP. Furthermore, due to concerns about the effects of motion, scans are acquired in stacks that must be reconstructed and motion corrected prior to analysis. Finally, T1 and T2 image resolution of the neonates is slightly reduced relative to the adults, but brains are much smaller resulting in reduced resolution and increased partial volume.

**Table 3:** Comparison between HCP and dHCP pipelines

|  | HCP | dHCP |
| --- | --- | --- |
| Image Resolution | $0.7mm^3$ | $0.8mm^3$ ($0.5mm^3$ after reconstruction) |
| Total scanning time | 4 hours | 76 minutes |
| **Preprocessing:** |  |  |
| Gradient Distortion Correction | Yes | No |
| Read-out distortion correction | Yes | No |
| Brain Extraction | Propagation of atlas mask | BET |
| T1 - T2 registration | BBR | Rigid Alignment, BBR |
| Bias Correction | $\sqrt{T1 * T2}$ | N4 |
| **Segmentation / Surface extraction** |  |  |
| Performed on | T1 | T2 |
| Tissue Segmentation | FreeSurfer | Draw-EM |
| White and Pial Surface Extraction | modified FreeSurfer | Schuh et al. (2017) |
| Surface Inflation | FreeSurfer | FreeSurfer re-implementation |
| Spherical Projection | FreeSurfer | spherical MDS |
| Myelin Mapping | T1/T2 | T1/T2 |

The HCP performs gradient and read-out distortion correction prior to estimation of surfaces. Gradient distortion, in particular, is motivated on account of the HCP’s use of a specially designed 3T Skyra scanning system, where the head is ~ 5 cm above the iso-centre of the scanner. This makes gradient non-linearities more pronounced for this system relative to those from standard 3T scanners. Read-out distortion correction is performed through collection of a field map. The dHCP has less noticeable distortions due to the use of spin-echo readouts in the dHCP acquisition. Additionally, distortions are less extreme in neonates due to differences in the air tissue interfaces from adults. As such the dHCP does not perform these corrections.

Surface extraction, within the HCP pipeline, is performed using a refined version of the FreeSurfer recon-all method Fischl et al. (1999b, c), which adjusts for the high spatial resolution of HCP data, and uses T2 surfaces to refine and improve the placement of white and pial surfaces. This accounts for limited contrast between grey and CSF tissue within adult T1 images. Modified pre-processing is performed prior to recon-all with brain tissue masks propagated (following non-linear registration) from the MNI Template, and T1-T2 registration is performed using a 6-DOF BBR Greve and Fischl (2009).

Image processing for the dHCP is performed on T2 images which provide better tissue intensity contrast in the neonatal age groups. Brain extraction is performed liberally on each subject’s T2 image using FSL BET Smith (2002), and this brain mask is then refined from the Draw-EM segmentation. T1-T2 registration is performed with an initial 6-DOF rigid registration of the T1 image to the T2 image based on WM and sub-cortical structures (i.e. excluding cGM and CSF), followed by BBR. Masks are then propagated from T2 to T1 image data through the computed transformation. The reduced resolution, and increased partial volume of neonatal data motivates use of specially designed tissue segmentation (Makropoulos et al., 2012, 2014, 2016) and surface extraction protocols Schuh et al. (2017) (Sections 3.2,3.3). Further differences between the dHCP and HCP pipelines, are motivated by a desire to reduce software overhead: therefore, surface inflation is performed using an in-house re-implementation of the FreeSurfer method, and spherical projection is performed using a specially designed spherical MDS tool (Section 3.4). Outside of the stated differences to the pre-processing and surface extraction pipelines, cortical features (cortical thickness, cortical surface curvature, sulcal depth, T1/T2 myelin maps) are estimated similarly to the HCP.

## 6. Discussion

This paper presents a fully automatic pipeline for brain tissue segmentation and cortical surface modelling of neonatal MRI. All methods have been tuned on images collected using the dHCP protocol, and take advantage of improvements in image quality (gained from advances in acquisition, reconstruction and motion correction (Cordero-Grande et al., 2017; Hughes et al., 2016; Kuklisova-Murgasova et al., 2012)) to offer topologically correct (genus-0) surface representations images acquired between 28 and 45 weeks PMA. Manual QC of the images and surfaces suggest that, in the vast majority of cases, surfaces correspond closely with tissue boundaries, improving upon results obtained with state-of-the-art tissue segmentation tools.

The goal of the dHCP project is to develop the first four-dimensional model of emerging functional and structural brain-connectivity. Accordingly, the dHCP is pushing the boundaries of structural, functional and diffusion image acquisition, to improve the spatial resolution of T1 and T2-weighted images, the angular resolution of diffusion MRI (dMRI) and the temporal resolution of functional MRI (fMRI), relative to previous neonatal and fetal studies.

The results in this paper represent the foundations of a surface-constrained set of functional and diffusion analysis pipelines, inspired by frameworks proposed by the HCP (Glasser et al., 2013; Van Essen et al., 2013), and based on FreeSurfer methods for surface extraction (Fischl et al., 1999b; Fischl, 2012). Studies have shown that surface-constrained analyses: improve the localisation of functional units along the cortical surface (Fischl et al., 2008; Glasser et al., 2013, 2016b; Van Essen et al., 2012); reduce the mixing of white and grey matter fMRI signals during smoothing (Glasser et al., 2013); and increase the alignment of functional areas during registration (Durrleman et al., 2009; Fischl et al., 1999c; Lombaert et al., 2013; Robinson et al., 2014; Wright et al., 2015; Yeo et al., 2010).

However, surface-constrained analyses rely on highly (geometrically) accurate reconstructions of the cortical surface. FreeSurfer-derived HCP pipelines do not work for neonates, whose brains are smaller and still developing. Neonatal images present inverted tissue intensity profiles, hypo/hyper intense white-matter patches and, due to the smaller size of the imaged structures, even for same image resolution, they are more prone to partial volume effects than for adult MRI. Further, dHCP neonates are imaged during natural sleep. As a result, motion artefacts are a concern, with a small subset of acquisitions heavily affected.

FreeSurfer methods for surface extraction depend on tissue segmentation models that are largely intensity-based (Dale et al., 1999). These are not robust to noise and are ill-suited to the inverted tissue intensity distributions, and increased partial volume of neonatal data. Further, FreeSurfer white cortical surface mesh modelling approaches rely on tessellating topologically correct white matter tissue segmentations, and FreeSurfer pial surface extraction tools require tissue intensity information; making implicit assumption that the image provided is a T1 (with light GM and dark CSF) (Dale et al., 1999).

These restrictions have required the development of bespoke methods for tissue segmentation and cortical surface mesh extraction for neonates and this paper summarises a refined pipeline for cortical surface mesh modelling and inflation that has been tuned on dHCP-acquired neonatal data. Within this pipeline, some tools, such as the Draw-EM tool Makropoulos et al. (2014), existed before the project. However, refinements have needed to be made to optimise this tool to work well with term-born neonates, and the increased resolution and contrast in the new cohort. This is because previous developmental brain studies performed by this project consortium, have been conducted predominantly on preterm data. The contrast and appearance of these images is, due to the increased CSF and fewer cortical folds, very different from healthy neonates at an equivalent developmental stage. Therefore, this has required significant improvements to the Draw-EM tool in order to: allow segmentation of hypo and hyper intense patches of white matter (within the broader class of all WM tissue); and improve alignment of tissue class priors in order to improve the accuracy of the segmentation, particularly for gyral crowns. Prior to these improvements it was common to observe segmentations missing broad sections of cortical folds.

Other tools have been newly implemented and improve on existing tools in the literature. These include the methods for white and pial surface mesh modelling (Schuh et al., 2017), which improves on the segmentation boundary produced using Draw-EM by introducing intensity-based terms to reduce the impact of partial volume. This approach is inspired by FreeSurfer (Dale et al., 1999), which also refines surfaces based on image intensity values. However, our neonatal approach is tuned to work with neonatal T2 intensity information, allowing extraction of white and pial surfaces, using novel deformable surface forces specifically designed for the reconstruction of the neonatal cortex. Surface QC suggests this method generates surfaces that agree well with tissue boundaries in 85% of cases, as assessed by expert raters.

Finally, methods for surface inflation and spherical projection, represent direct re-implementation of the inflation approach used in FreeSurfer (Fischl et al., 1999b) and the spherical MDS embedding approach proposed by (Elad et al., 2005). Their inclusion within the pipeline is therefore designed to reduce software overhead. Note, all tools described in this paper are open source and are available as part of MIRTK^4^. The full pipeline is available as open-source ^5^ and this paper accompanies the first open data release ^6^ (version 1.1). The pipeline has been tested with different Linux distributions and macOS, and can be executed on a standard PC with 8 GB RAM. A typical dataset can be processed in around 18 hours using 8 CPU cores (segmentation: 7 hours, surface processing: 10 hours, additional processing: 1 hour).

It is important to stress that surface extraction techniques described in this paper are not infallible. Manual QC suggests that in a minority (2%) of cases (typically overlapping with highly motion data sets) entire folds may be excluded from the automatic segmentation. These errors are propagated to the surface extraction, which despite accounting for tissue intensity may not be able to recover from major segmentation errors, due to the significant impact of partial volume. Surface QC suggests there exists regional biases, highlighting the temporal and occipital lobe as regions most effected. For these regions, in particular, comparisons of morphological features such as cortical thickness, must be interpreted with caution.

The methods described in this paper are by no means designed to represent the breadth of tools available for neonatal image processing. Indeed, several alternative methods for neonatal segmentation have been proposed including (Xue et al., 2007; Hill et al., 2010; Li et al., 2012, 2014a, 2016; Hazlett et al., 2017). Different techniques have also been proposed to ensure consistent segmentation/reconstructed surfaces between different time points in longitudinally acquired data (Nie et al., 2012; Dai et al., 2013; Wang et al., 2013; Li et al., 2014a, c) is not something that has been explicitly dealt with in our pipeline as relatively few data sets (8%) are longitudinal. Further, longitudinal deformation studies that have been performed using this data indicate that the patterns of deformation observed are biologically plausible Robinson et al. (2017). Nevertheless, this type of approach may prove more valuable when the project expands to fetal data.

In this paper, we extend upon our previous segmentation algorithms Makropoulos et al. (2012, 2014). These methods are state-of-the-art generating the most accurate results on a recent neonatal segmentation challenge (Isgum et al., 2015), and demonstrating robustness with respect to segmentation of the neonatal brain at different scan ages, despite large differences in appearance (Makropoulos et al., 2014, 2016). Part of the reason for the success of these methods is the inclusion of terms that reflect prior knowledge derived from manually annotated tissue atlases (Serag et al., 2012), and knowledge of how partial volume reflects tissue contrast. Nevertheless, in very low contrast areas, even our revised automated segmentation may still fail. In these cases, the deformable surface model may also not recover the correct boundaries, as information provided by tissue intensities is obscured through partial volume. Complete correction of these issues will likely require development of completely new tools for segmentation and surface extraction that incorporate enhanced prior information. This might be inferred from higher resolution images, or improved manual segmentations.

The dHCP is not the only large-scale project focused on open-release of developing brain data. The baby connectome project, run as a collaboration between the University of North Carolina and the University of Minnesota, seeks to provide data on how the human brain develops from birth through early childhood by collecting 500 data sets of infants between 0 and 5 years of age. Half of this data will be collected as longitudinal scans. The data set does not overlap with the dHCP, where neonatal scans are all collected before and within a few days of birth, but instead will collect valuable information on how the brain develops in the years after birth. An important future goal of both projects will be the provision of tools to link these data sets.

In the absence of ground truth knowledge of the correct tissue class membership and surface geometry of all neonatal datasets, assessment of different parts of the proposed pipeline has been performed through quality control inspection of the images, resulting tissue segmentations and surface reconstruction by two expert raters. Quality was assessed through metrics designed to meet our assessment of the needs of the community, including evaluation of the amount of motion, and anatomical correctness of the segmentation and surfaces. Results suggest that, out of 160 images, 2% had to be discarded on account of the effects of motion being too severe to be corrected during reconstruction. Of the remaining data sets, only an additional 1% had to be discarded due to poor segmentations. Surface reconstruction quality was assessed on 43 subjects for 20 ROIs each (860 ROIs) with only 2% of the ROIs presenting poor quality.

There are limitations to this QC in as much as evaluation was performed manually, by a relatively small number of raters. In future we aim to train a classifier that will automatically score an image, segmentation or surface based on the existing manual scores. Nevertheless, quality assessment of neurological data is extremely challenging on account of the lack of ground truth. We release the data openly and encourage users to feedback their experience.

Structural image data for the dHCP have been collected at 0.8 *mm*^3^ resolution, and reconstructed at 0.5 *mm*^3^ isotropic resolution, with the goal of ensuring the best possible quality surface extraction. We recommend that any study, seeking to replicate results from the dHCP pipeline, acquire scans at a similar, or higher, resolution. Nevertheless, the methods used in the paper are robust to the increased partial-volume effects of low-resolution neonatal MRI data. In particular, the Draw-EM method is robust to lower-resolution data, having been initially developed for data acquired from overlapping stacks at 0.86 × 0.86 mm in-plane resolution, slice thickness 2 mm, and overlap of 1mm (Makropoulos et al., 2012). In principle, as surface extraction is contingent upon the result of successful segmentation, surfaces could be extracted from low resolution data. However, the parameters of intensity-based surface correction tools used within the proposed pipeline would likely need to be re-tuned, and it is possible that the proportion of errors, such as missing gyri, would be increased. Note, for any study seeking to apply proposed pipeline to new data, we must stress that the parameters used in the paper have not been quantitatively assessed. Optimal choice of parameters is not feasible due to the lack of available ground truth. Subjective choices have been made that balance prior understanding, that the cortical surface is smooth, against a perfect fit with tissue contrast (which is impacted by noise and tessellation of the images.

The surface meshes generated through this pipeline will serve as a basis from which structural, functional and diffusion datasets will be compared across populations and over time. Comparisons between data sets will be facilitated through development of standard “grayordinate” CIFTI template spaces Glasser et al. (2013), which will allow compact representations of highdimensional dMRI and fMRI data. Unlike for adult data these will evolve spatio-temporally to account for the rapid development of neonatal brains. Methods for generating these atlases are considered out of the scope of this paper, but interested readers should refer to the frameworks described in Schuh et al. (2015) and Bozek et al. (2016b, 2017) for more details. Alternative techniques have also been proposed for construction of perinatal surface atlases (Hill et al., 2010; Li et al., 2015c).

Within this paper we present cortical T1/T2 myelin maps using the method proposed in Glasser and Van Essen (2011) and adapted for use in neonates as shown in Bozek et al. (2016a). The use of HCP-standard cortical myelin maps is chosen to allow comparison with adult HCP data, and also on account the choice of scans available through the dHCP protocol. Within this limitation, it is important to note that the neonatal myelin maps were generated from a IR-TSE T1 sequence, rather than the MPRAGE as used in Glasser and Van Essen (2011). Outside of these restrictions, it is important to acknowledge that myelin has been studied long before in the past (Barkovich et al., 1988; Counsell et al., 2002; Prastawa et al., 2005). Furthermore, for studies specifically interested in the development of cortical myelin, there exist more advanced methods for sensitising MRI acquisition to myelin content (Borich et al., 2013; Deoni et al., 2011; Dinse et al., 2015; Laule et al., 2008; MacKay and Laule, 2016; Melbourne et al., 2013; Stuber et al., 2014). Some of these in particular, have been shown to be sensitive to developmental changes in myelin (predominantly of the white matter) (Deoni et al., 2011, 2015). A more thorough review of methods for myelin mapping is provided in MacKay and Laule (2016).

In this work we provide an example cortical parcellation based on the neonatal atlas constructed by Gousias et al. (2012), which is incorporated within the Draw-EM segmentation tool. The regions in the Gousias et al. (2012) atlas have been specifically annotated for the neonatal brain and are different from the Desikan-Killiany atlas (Desikan et al., 2006) that has been developed for adults, and is being used in FreeSurfer. Figure 25 presents an example cortical surface parcellated with the two different atlases. Alternative multi-label atlases could be used for the segmentation and cortical parcellation (de Macedo Rodrigues et al., 2015; Alexander et al., 2016), however these are not publicly available and could not be included in our publicly available pipeline software. The Melbourne Children’s Regional Infant Brain (M-CRIB) atlas constructed by Alexander et al. (2016) would be particularly interesting as it replicates the Desikan-Killiany protocol and delineates 100 regions in the T2 scans of 10 term-born neonates. This could help to link regions between the perinatal and adult period. In future, it may also prove valuable to delineate the cortex based on patterns of structural or functional connectivity, using methods such as Arslan et al. (2015); Craddock et al. (2012); Gordon et al. (2014); Parisot et al. (2016a, b), or train a classifier that will propagate the Glasser et al. (2016a) method for multi-modal parcellation of the adult cortex onto neonatal brains.

**Figure 25:**
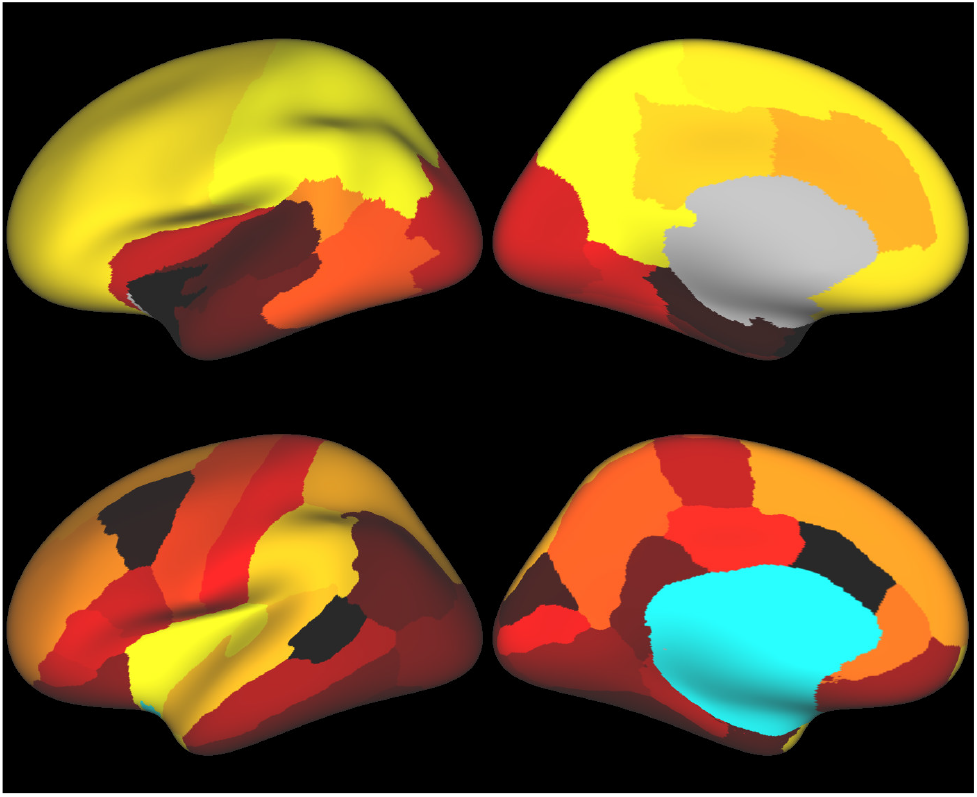
Comparison of the Draw-EM labels, based on Gousias et al. (2012), (top) and FreeSurfer labels, based on Desikan et al. (2006), registered and transformed on an example subject (bottom). Note that the colours used for the different atlas regions are independently drawn for the atlases, and not correspond between the two atlas labellings.

Most importantly, application of the developed methods will allow to study the developing brain and deviations due to pathology. Numerous works have been published in the literature that have improved our understanding of the perinatal brain (Ball et al., 2013b, 2017; Counsell et al., 2013; Doria et al., 2010; Kapellou et al., 2006; Keunen et al., 2017; Krishnan et al., 2017; Xue et al., 2007; Dubois et al., 2008a, b; Pienaar et al., 2008; Rodriguez-Carranza et al., 2008; Awate et al., 2010; Hill et al., 2010; Rathbone et al., 2011; Moeskops et al., 2013; Li et al., 2014b, a; Meng et al., 2014; Nie et al., 2014; Wang et al., 2014; Engelhardt et al., 2015; Lefevre et al., 2015; Li et al., 2015b; Moeskops et al., 2015; Li et al., 2016; Hazlett et al., 2017). Studying the developing connectome will open up many opportunities in future, not least because neonatal brain imaging data is changing rapidly, at scales that can be clearly resolved using current MRI technology. Work is already under-way for the reconstruction, artefact-removal and surface-projection of neonatal functional, and diffusion data sets, and the extension of all pipelines to fetal cohorts. Together, these image sets (and the genetic, behavioural and clinical information that support them) will allow improved understanding of the spatio-temporal development of the cortex at the millimetre scale. They will enable modelling of the mechanisms of cognitive development, and potentially allow to further link the perinatal brain development to the childhood and adult period. Analysis of these data sets will provide a vital basis of comparison from which preterm development, and the causes of neurological conditions such as cerebral palsy or autism, may become better understood.

## Acknowledgements

The research leading to these results has received funding from the European Research Council under the European Unions Seventh Framework Programme (FP/2007-2013) / ERC Grant Agreement no. 319456. We are grate-ful to the families who generously supported this trial. The work was supported by the NIHR Biomedical Research Centers at Guys and St. Thomas NHS Trust.

## Appendix A.

**Figure A.1:**
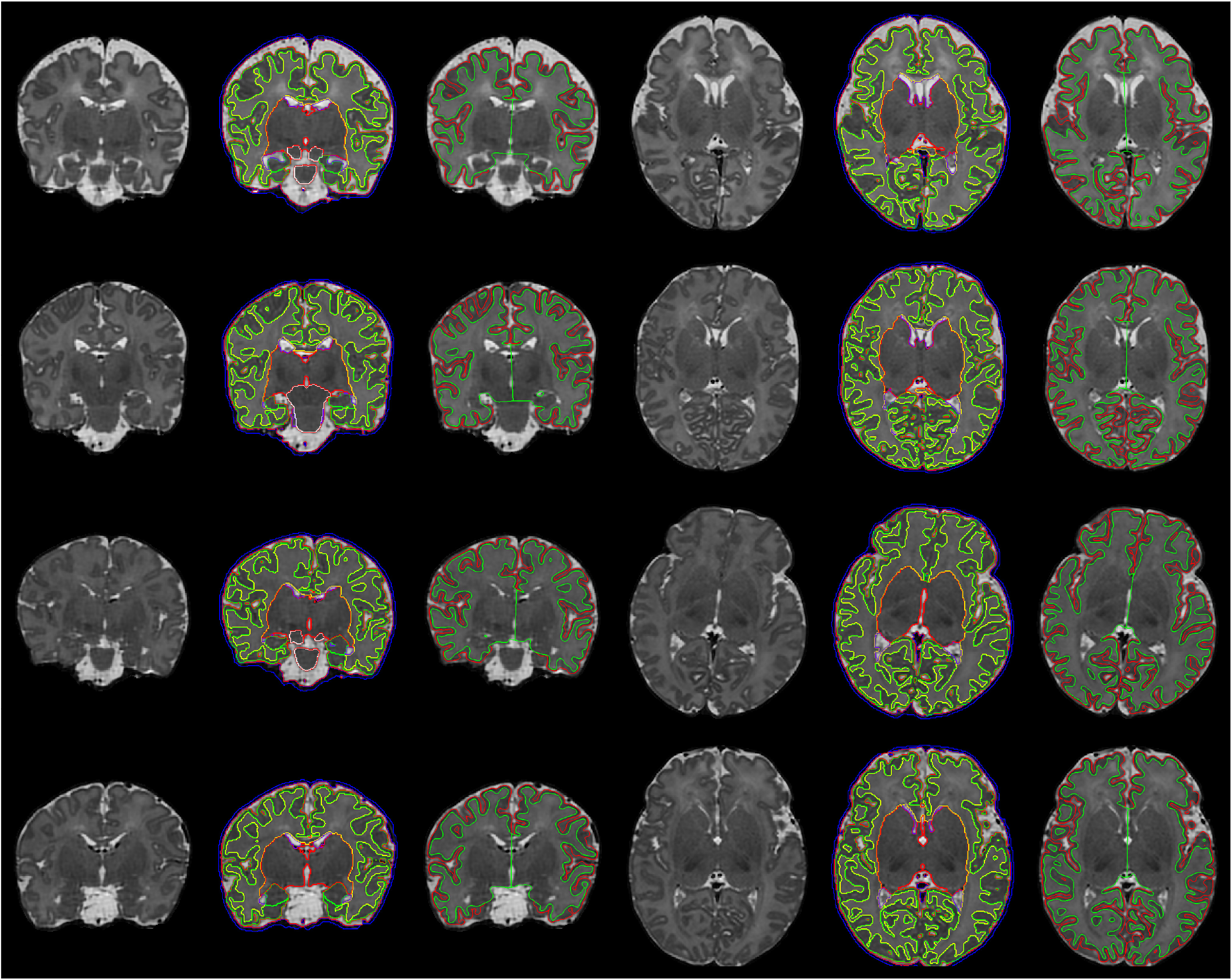
Example segmentations and surface reconstructions of subjects scanned at 37, 38, 39 and 40 weeks (from left to right: coronal T2, coronal T2 with segmentation overlaid, coronal T2 with white and pial surfaces overlaid, axial T2, axial T2 with segmentation overlaid, axial T2 with white and pial surfaces overlaid).

**Figure A.2:**
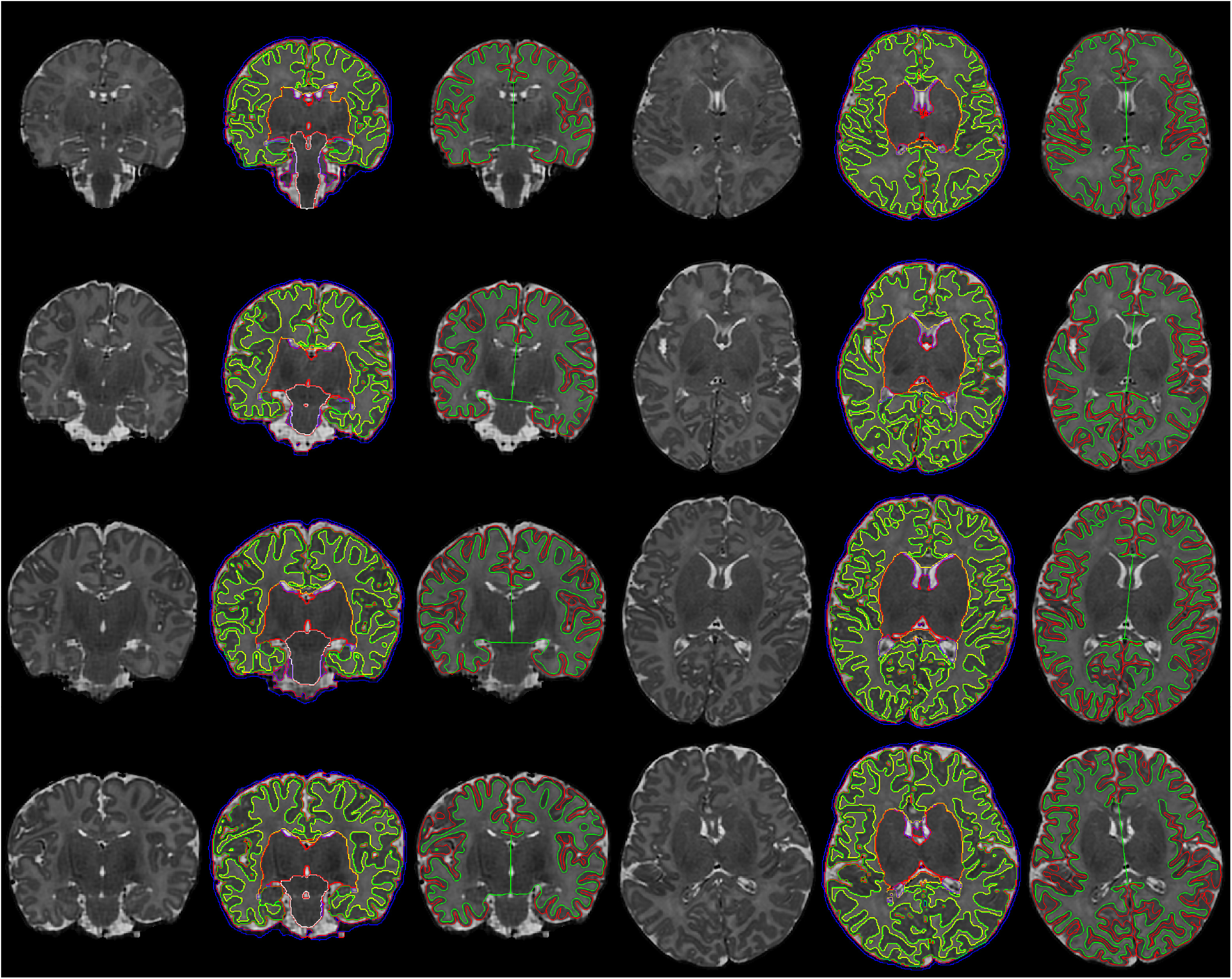
Example segmentations and surface reconstructions of subjects scanned at 41, 42, 43 and 44 weeks (from left to right: coronal T2, coronal T2 with segmentation overlaid, coronal T2 with white and pial surfaces overlaid, axial T2, axial T2 with segmentation overlaid, axial T2 with white and pial surfaces overlaid).

**Figure A.3:**
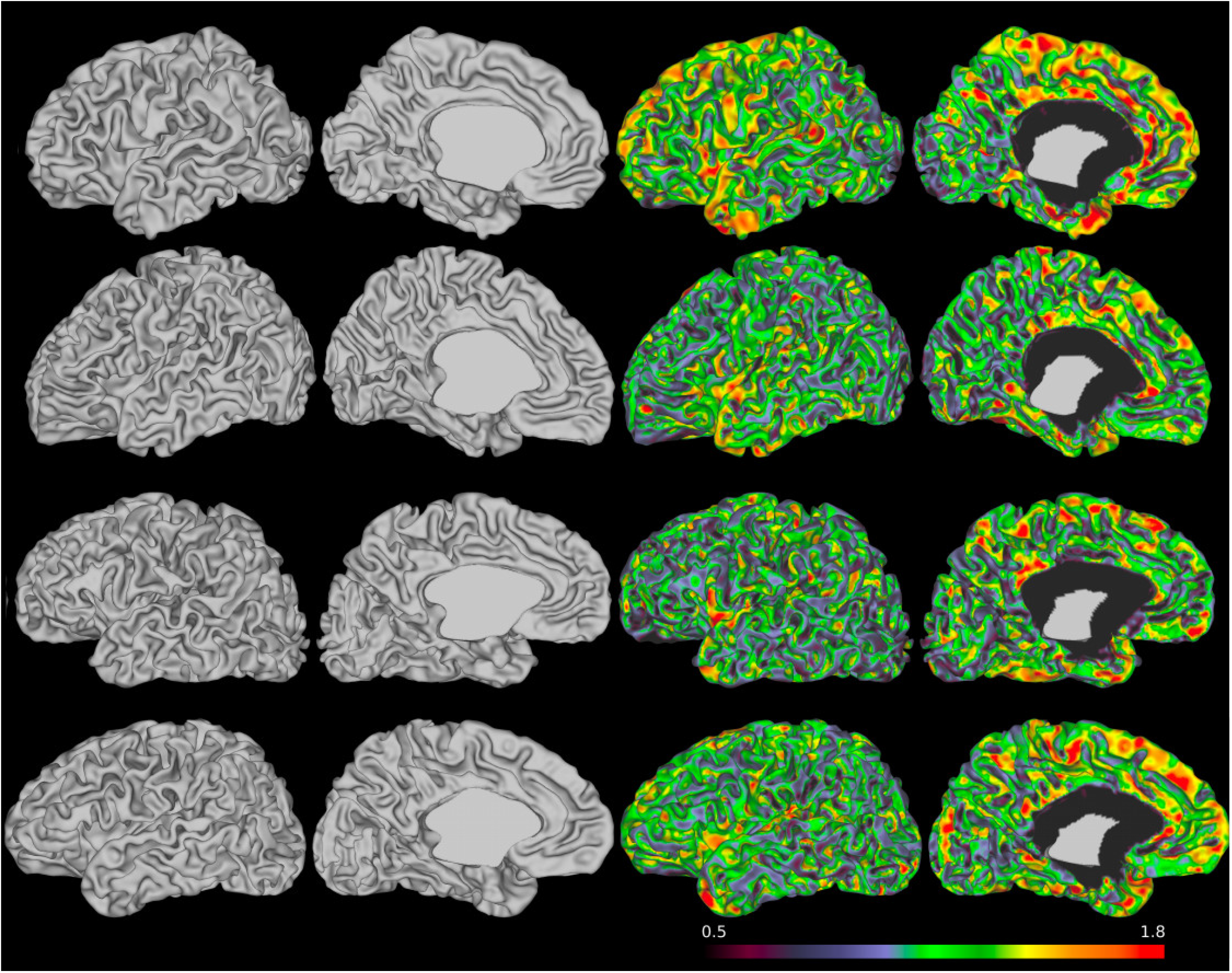
Example left hemispheres white surface reconstructions of subjects (same as Fig. A.1) scanned at 37, 38, 39 and 40 weeks with cortical thickness overlaid (from left to right: lateral view of white surface, medial view of white surface, lateral view of white surface showing cortical thickness, medial view of white surface with cortical thickness).

**Figure A.4:**
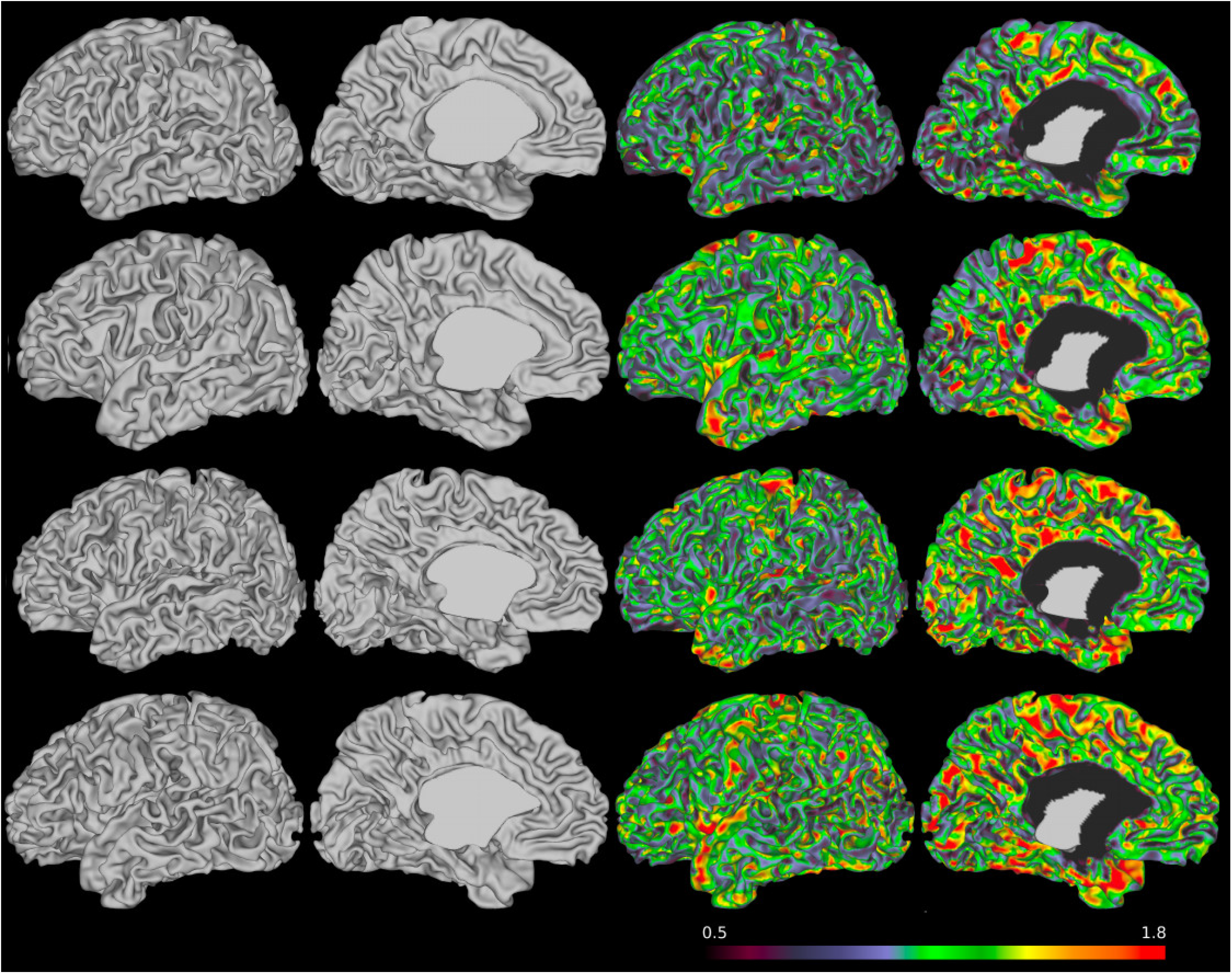
Example left hemispheres white surface reconstructions of subjects (same as Fig. A.2) scanned at 41, 42, 43 and 44 weeks with cortical thickness overlaid (from left to right: lateral view of white surface, medial view of white surface, lateral view of white surface showing cortical thickness, medial view of white surface with cortical thickness).

**Figure A.5:**
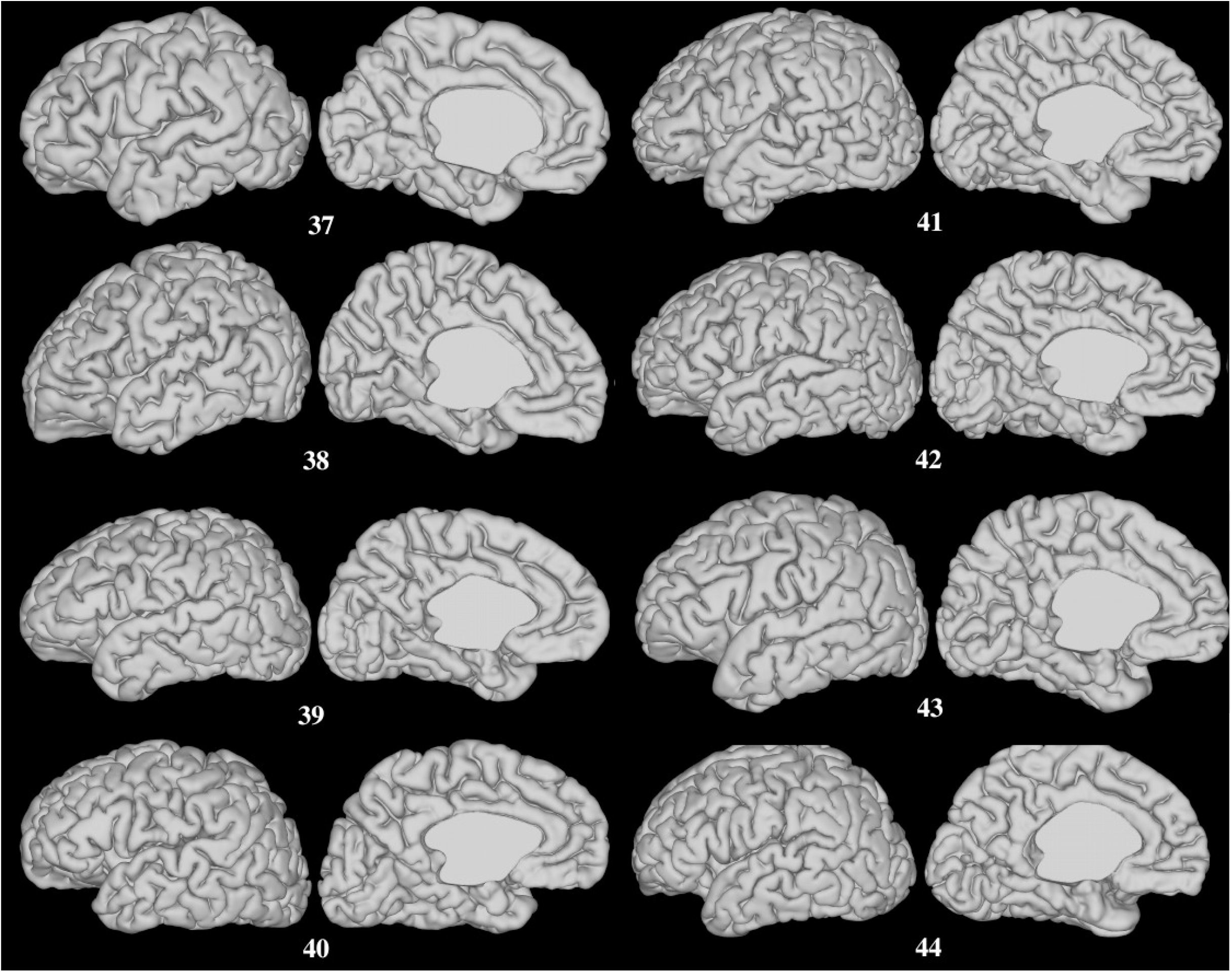
Example left hemisphere pial surfaces for all subjects in Figs. A.3 and A.4). Lateral and medial views are shown

## Footnotes

2 http://brain-development.org/brain-atlases/neonatal-brain-atlas-albert

3 https://github.com/MIRTK/DrawEM

4 https://biomedia.doc.ic.ac.uk/software/mirtk/

5 https://github.com/DevelopingHCP/structural-pipeline

6 https://data.developingconnectome.org

